# A battery of image classification challenges reveals shared and distinct object categorization behavior across monkeys, humans, and deep networks

**DOI:** 10.1101/2025.04.02.646407

**Authors:** Han Zhang, Zhihao Zheng, Jiaqi Hu, Qiao Wang, Mengya Xu, Zhaojiayi Zhou, Zixuan Li, Gouki Okazawa

## Abstract

Humans categorize objects at multiple levels of abstraction—animate versus inanimate, big versus small, and many other attributes. Despite its apparent challenge, the advent of deep neural networks (DNNs) has demonstrated that complex visual processing alone can support such classification without language or human-specific knowledge. This raises a natural question: to what extent can non-human primates, without language, perform such categorization? Although basic object-recognition behavior in monkeys such as similarity judgment has been extensively studied, their ability to classify objects across diverse rules remains poorly characterized. Here, we developed a task paradigm that enabled us to train monkeys on a large battery of binary classification tasks using natural object images, spanning more than 10 rules, such as animate versus inanimate, natural versus man-made objects, and mammalian versus non-mammalian animals. We then evaluated their performance on novel, generalization stimuli. Monkeys acquired each rule in a few days, generalized it well, and exhibited error patterns consistent with human judgments. At the same time, their classification performance correlated more strongly with that of visual DNNs trained without language input, whereas human performance was better explained by language-informed DNNs. These results provide an important benchmark for the capacity of biological neural networks to perform image classification without language and human-specific knowledge.

## Introduction

Humans can readily judge whether a visually perceived object belongs to various categories across multiple levels of abstraction, such as animate, mammal, big, and so on. Earlier theories proposed that basic-level categories (e.g., “dog” and “cat”) are classifiable because exemplars share visual and other features (Rosch et al. 1976), whereas more abstract categories (i.e., superordinate categories; e.g., animate) tend to lack defining features (Murphy and Medin 1985; Rosch et al. 1976) and rely more on semantic knowledge and experience (Lambon Ralph et al. 2017; Murphy 2004). However, the rise of deep neural network models (DNNs) has transformed our view of how apparently complex object-classification tasks can be solved by sophisticated visual processing alone. Since then, many studies have compared internal object representations between DNNs and biological brains (Khaligh-Razavi and Kriegeskorte 2014). In particular, research in macaque monkeys has demonstrated strong similarities between ventral-pathway neural activity and DNNs (Yamins et al. 2014), reinforcing the notion that visual learning and processing underlying core object recognition are shared across humans and animals (Kar and DiCarlo 2024).

However, surprisingly little is known about whether, and to what extent, monkeys can classify objects into a variety of high-level categories as humans do, because most previous behavioral studies focused on basic object-recognition performance, such as similarity judgment (Koopman et al. 2017; Rajalingham et al. 2015). For example, Rajalingham et al. (2015) systematically compared human and monkey behavior using a match-to-sample task with 3D object stimuli. With this design, they were able to measure monkeys’ confusion rates across numerous object identities, revealing a striking similarity to human performance. But this approach does not readily scale to higher-level categorization, as the task design cannot easily prompt animals to attend to more abstract features; indeed, the observed confusion patterns did not appear to clearly reflect superordinate structures such as animacy (Rajalingham et al. 2015; but see Cherian et al. 2025). In contrast, earlier psychological studies in humans have demonstrated profound influences of language and conceptual knowledge on object categorization (Murphy 2004). Thus, critical gaps between humans and monkeys may emerge when they are required to report higher-level categories. This question is timely, as growing efforts are being made to develop DNNs whose representations of abstract object concepts align with those of humans (Du et al. 2025; Muttenthaler et al. 2025; Vong et al. 2024).

Prior to the proliferation of DNNs, multiple studies actually challenged monkeys with various image classification tasks, but they had their own limitations. For example, a series of studies by Fabre-Thorpe et al. (Delorme et al. 2000; Fabre-Thorpe 2003; Fabre-Thorpe et al. 1998; Fize et al. 2011) demonstrated that monkeys can report the presence or absence of an animal in rapidly presented natural photographs using a Go/No-Go paradigm. Other studies have shown that monkeys can classify trees vs. non-trees (Vogels 1999), humans vs. non-humans (Schrier and Brady 1987), monkeys vs. non-monkeys (Yoshikubo 1985), food vs. non-food (Fabre-Thorpe et al. 1998; Santos et al. 2001), cats vs. dogs (Eldridge et al. 2018; Freedman et al. 2001; Minamimoto et al. 2010), and cars vs. trucks (Minamimoto et al. 2010). But each study typically tested only one or, at most, two classification rules, mostly using basic-level categories, and to our knowledge, there has been no systematic quantification of which classification rules monkeys can learn and which they cannot. Moreover, it was obviously impossible for earlier studies to compare monkey behavior with DNNs. Therefore, a systematic characterization of monkey behavior is still much needed; yet classification tasks are challenging, as monkeys must be trained on each rule individually, and performing a test battery becomes impractical if training for each task requires extended durations.

Here, we overcome these challenges by devising a novel binary decision-making paradigm that can train monkeys quickly, allowing us to test a series of image classification challenges with three monkeys (more than 10 tasks, 26 including subtasks, *∼*315K trials total; see Movies 1-3 for examples). The monkeys learned many classifications, such as animate vs. inanimate, natural vs. artificial, mammal vs. non-mammal, big vs. small, and their performance was correlated with that of humans. At the same time, they failed to learn some more abstract rules that humans could readily acquire. Overall, their behavior was better explained by purely visual DNNs without language input, whereas human performance was better explained by language-guided DNNs. While our paradigm does not answer whether monkeys internally represent these concepts (e.g., “mammal”), their performance provides an important benchmark for the extent to which biological neural networks can succeed in image-classification challenges without access to language and human-specific knowledge.

## Results

### Monkeys quickly learn many visual classification tasks defined at various levels of abstraction

Our task is a version of the binary decision-making paradigm, in which monkeys categorize an object image into one of two options according to a hidden rule (Fig. 1A). The experiments were conducted using a touchscreen system installed on the monkey’s home cage (Supplementary Fig. 1A). During each trial, after the monkey touched a red dot, an object image appeared together with two gray target boxes. The monkey had to touch the object image and drag it to one of the two target boxes. The image position moved along with the monkey’s finger until it reached a box. Upon reaching, the two boxes disappeared, and the same object image was revealed under the correct box, together with a juice reward for the correct choice. We reasoned that direct interaction with an object image encouraged monkeys to rely on the object’s properties to make a behavioral choice. Furthermore, this dragging motion required monkeys to slow down their choice reports (*>* 0.5 sec to reach a target), potentially suppressing random impulsive choices and allowing them to make more deliberate decisions (Kaufman et al. 2015). The categorization rule remained the same within a session, and the same rule continued across sessions until the monkey’s performance plateaued. The monkeys had to infer this rule based on the feedback they received on each trial.

This task design enabled rapid, high-throughput training of monkeys. We first trained this task structure using amorphous shape stimuli for a month (Supplementary Fig. 1B, C) and then started the first object classification experiment: animate versus inanimate classification (Fig. 1B-F). The training images consisted of a variety of grayscale images without background (Fig. 1B), but all three tested monkeys achieved high classification performance within a few days of training (Fig. 1C-D; averaging 760 trials per day). They started to show above-chance performance from Day 1 (*p <* 0.011 for all monkeys, binomial test), and their correct rate monotonically improved over the days, plateauing at approximately Days 4-6 with a performance of *∼*90% (Fig. 1D). The same training images (*n* = 96) were repeatedly presented during these sessions, but the monkeys started to show above-chance performance only after a few repetitions per image (Fig. 1D inset; *p <* 0.05 at 3, 4, and 9 repetitions for monkeys Ol, Ju, and El, respectively, binomial test).

**Figure 1:**
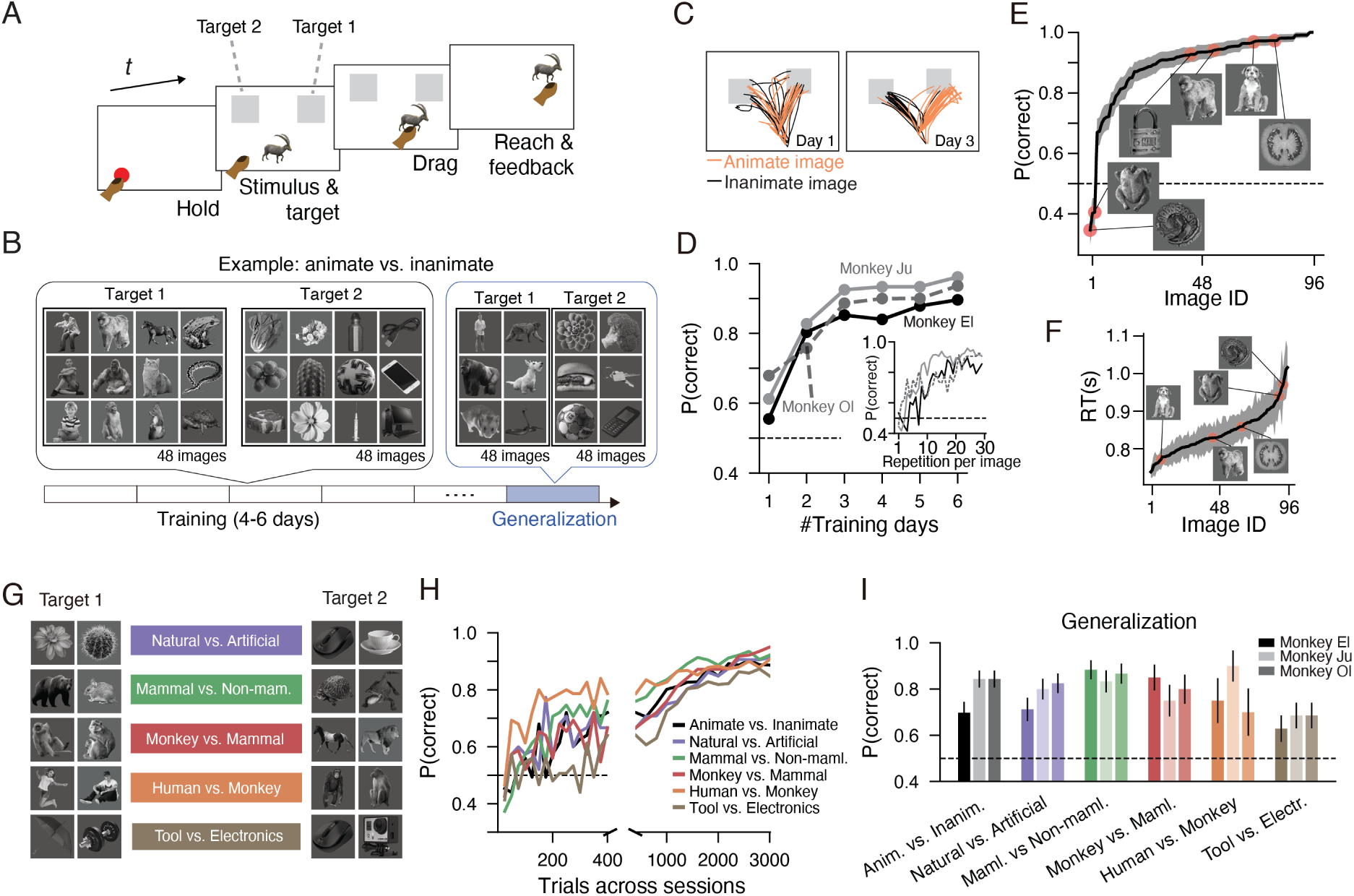
Monkeys rapidly learn various abstract object concept classification tasks. (**A**) Object drag task. In each trial, after the monkey held a fixation dot, an object image and two gray target boxes appeared. The monkey had to touch and drag the image to the correct box to receive a reward. Two boxes were associated with different object concepts. (**B**) We used grayscale photographs of objects with no background in this first set of experiments. We used ∼100 training images to train monkeys and measured their generalization performance with test images. (**C**) Example reach trajectories showed that monkeys tended to choose targets randomly on Day 1, but by Day 3, they chose correct targets in many trials. (**D**) All monkeys’ correct rates rapidly improved during the first three days of the animate vs. inanimate task, reaching 85-90% and then plateauing (chance level: 50%). Before the performance plateaued, monkeys saw each training image 10-20 times (inset). (**E**) After learning of the animate vs. inanimate task (post-Day 3), monkeys made more than 90% correct choices for many images, but performed poorly on certain images. The plot is the average of the three monkeys, with images ordered by correct rates. The shading indicates S.E.M. across trials. (**F**) Reaction times (RTs), measured as the time from stimulus onset to reaching a target, also varied across images. The plot is the average of the three monkeys, with images ordered by RTs. (**G**) We tested a variety of classification rules in succession. Each task used 60-96 grayscale training images. (**H**) Correct rates as a function of trial counts for each task. The data are averaged across the three monkeys. Left, 25-trial bin. Right, 200-trial bin. (**I**) After training on each task, the monkeys could generalize the rule to new stimuli they saw for the first time. The error bars represent S.E.M. across images. All images were used either under the CC0 license or the Pixabay content license.

Although their overall performance did not reach a 100% correct rate in six days, we found that they achieved a nearly 100 % correct rate for some images and performed poorly for some other images (Fig. 1E). Images with a nearly 100% correct rate were often prototypical examples of animate and inanimate categories such as monkeys, humans, and key locks, while images with poor correct rates tended to be atypical examples, such as a snake in a coil or a roasted chicken (defined as inanimate food). The patterns of correct rates were also largely consistent across monkeys (average of three comparisons: *R* = 0.49, *p <* 2.1 *×* 10^−4^ for all comparisons). The reaction times (RTs) also tended to be longer for these stimuli (Fig. 1F), and the drag trajectories for these stimuli occasionally showed patterns that resembled animals changing their minds (Supplementary Fig. 2; Kaufman et al. 2015).

We then confirmed that the monkeys could generalize the rules that they learned to new images seen for the first time (Fig. 1I). After the learning sessions, we introduced a new set of images (*n* = 96) that largely overlapped basic-level categories with the learned images but had different appearances (Fig. 1B). The monkeys’ correct rates for these new images were high from the first exposure (*p <* 6.6 *×* 10^−5^, binomial test).

Encouraged by these results, we tested a variety of classification rules based on both superordinate and basic-level object concepts (Fig. 1G). These tasks were natural (plants, food) vs. artificial (man-made) objects, mammal vs. non-mammal animals (reptiles/amphibians), monkeys vs. mammals, humans vs. monkeys, and tools vs. electronics. The monkeys also had no difficulty learning these tasks in a few days (Fig. 1H; *p <* 10^−10^ for all monkeys in all tasks on Day 4, binomial test) and generalizing the rule to new images that they had never seen before (Fig. 1I; *p <* 6.1 *×* 10^−6^ in all tasks aggregated across monkeys, binomial test). When we switched to the next rule, we did not give the monkeys any explicit cues, but they learned to adopt the new rule quickly.

### Monkey behavior depends on combinations of visual/categorical features

How did monkeys perform these categorizations? There are a couple of trivial hypotheses that have to be excluded. For example, monkeys might have simply formed associations between individual images and choices, and their choices for new stimuli in the generalization set (Fig. 1I) might be based on similar stimuli they had learned before. Alternatively, monkeys might have just learned a specific feature useful for categorization, such as the presence of faces or legs in the animate vs. inanimate task. Below, we show several control experiments in our test battery that excluded these possibilities.

One test involved a larger number of generalization images whose object categories were the same or different from those shown during training. We used images from the THINGS database (Hebart et al. 2019; Stoinski et al. 2024) and selected a subset of the concrete object concepts in the database (e.g., dog, spider, lemon, and table) to create three tasks (Fig. 2A): animate vs. inanimate (373 object categories, 3,683 images in total), natural vs. artificial objects (230 object categories, 1,915 images in total), and mammal vs. non-mammal animals (143 object categories, 1,668 images in total). For each task, we selected 100 images as a training set and then introduced the remaining images to test generalization. Importantly, the generalization images, which were shown only once, could be from object categories that were shown during training (“old”; Fig. 2A, second row) or were never shown (“new”; Fig. 2A, bottom two rows). This enabled us to test whether the generalization performance depended on the similarity to the trained images (i.e., whether the “old” categories showed greater performance).

**Figure 2:**
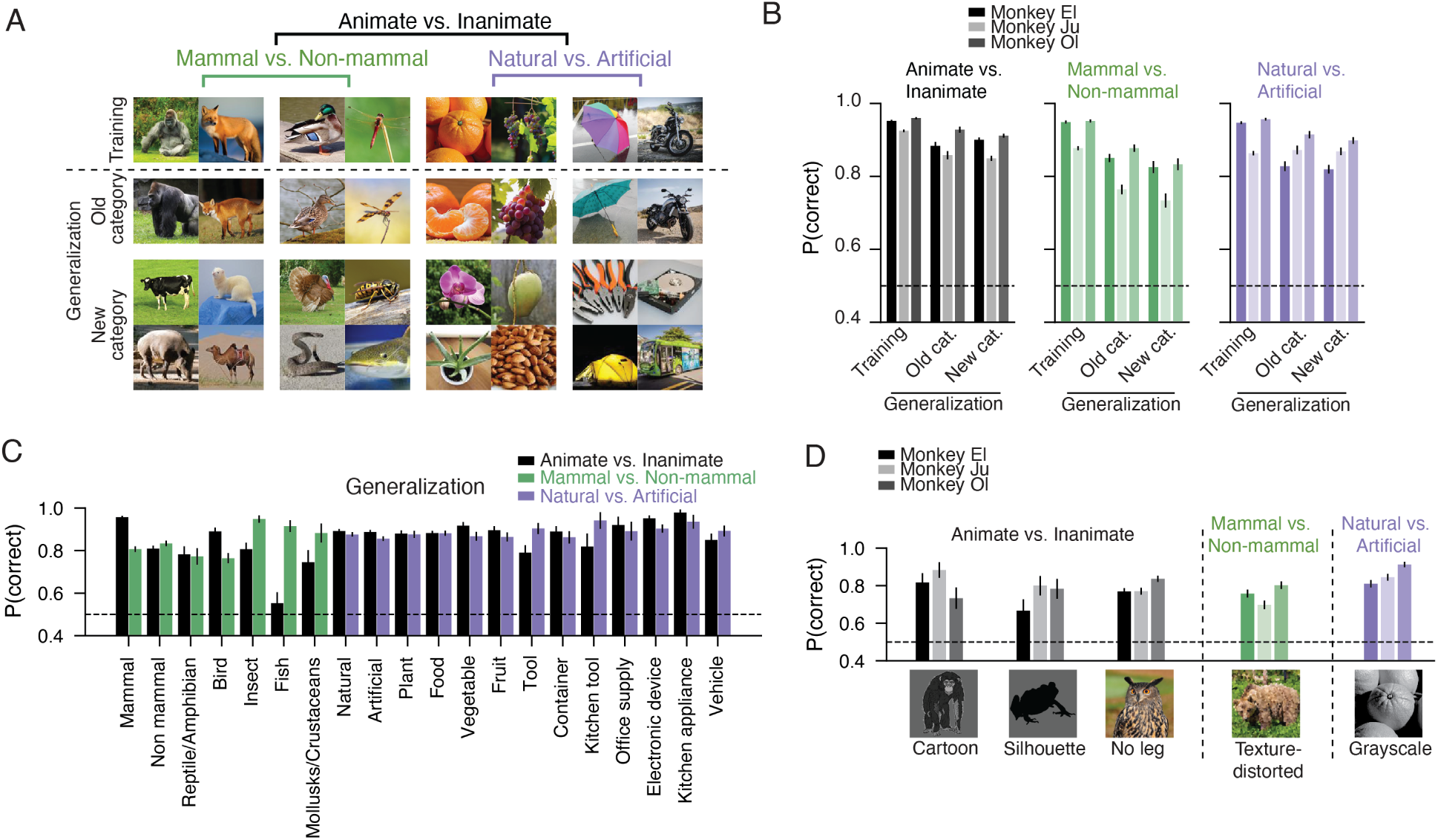
Behavior cannot be explained by exemplar-based strategies or some accidental features. (**A**) We assessed monkey performance in three tasks using large-scale image sets from the THINGS database (Hebart et al. 2019). We used 100 images to train monkeys (top row) and then presented 1,500-3,600 generalization images, containing “old” object categories present in training images (second row) and “new” categories that were never shown (bottom two rows). Each task used different training images. (**B**) All monkeys performed well on generalization images (75-93%) for both the “old” and “new” object categories. (**C**) Performance was generally high across a range of object categories. (**D**) Monkeys also performed well on control images that lacked certain visual features, such as cartoons without naturalistic textures, silhouettes without internal structures such as faces, and grayscale images without color. (**B-D**) Error bars indicate S.E.M. across object categories. All images were used either under CC0 license or Pixabay content license.

The monkeys performed these tasks without difficulty, despite experiencing much greater variability in image appearance than that in the previous image sets (Fig. 2B; Movies 1-3; *p <* 10^−10^ for all monkeys in the three tasks, binomial test), and importantly, their generalization performance to “new” object categories was largely comparable to that for the novel images of the “old” object categories they learned during training (Fig. 2B; *U* = 2, 360 *−* 14, 902, *p* = 0.086 *−* 0.98 across monkeys and tasks, two-tailed Mann-Whitney *U*-test, BF_01_ *>* 3.7 except for the mammal vs. nonmammal in monkey Ol, whose BF_01_ was 1.9). Their performances were also largely similar across a range of object categories in the datasets (Fig. 2C), except for “fish” images during the animate vs. inanimate task, likely because these images were less typical as animate (see Fig. 6 for the effect of image typicality). Thus, it is unlikely that the monkeys generalized the rules by judging the similarity to specific exemplars they learned during training.

To test if the monkeys relied on a particular feature in the images (e.g., face, leg-like shape, certain colors, textures), we also presented control images that lacked these features (Fig. 2D). In the animate vs. inanimate task, we presented cartoon images that lacked naturalistic textures, and silhouette images that retained only the outlines of objects without internal structures such as faces. In the THINGS image set, there were close-up images of animals without legs. The monkeys performed well on these images (Fig. 2D; *p <* 6.7 *×* 10^−3^ for all monkeys, binomial test). In the natural vs. artificial task, color could have been a strong cue (natural objects tend to be greenish), but we found that the monkeys performed well on grayscale images (81-91 % correct, *p <* 10^−10^ for all monkeys, binomial test). Finally, fur texture can be diagnostic for the mammal vs. non-mammal task, but the monkeys performed the task well even after distorting texture (see Methods; 70-80 % correct, *p <* 10^−10^ for all monkeys, binomial test). Thus, they likely did not rely on a single specific feature to solve the tasks.

To further examine whether the monkeys relied on intermediate-level image features (Freeman et al. 2013; Okazawa et al. 2015; Wang and Ponce 2026), we used “texform” images (Long et al. 2018)—synthetic images generated from noise by locally matching textural image statistics with natural images (Fig. 3A). These images lack higher-level image features but still preserve information diagnostic of object animacy (Long et al. 2018). If the monkeys’ classification relied purely on such intermediate-level features, they should be able to generalize their classification rule to texform images. The results did not support this; after fully learning the animate vs. inanimate task with natural images, the monkeys performed at chance level on the texform version of the task (Fig. 3B; *p* = 0.26 *−* 0.82, BF_01_ = 4.7 *−* 6.3, binomial test). When the monkeys were further trained on the texform images, they could learn the classification, but at a slower rate than that in the natural image categorization (Fig. 3C; *p <* 9.5 *×* 10^−4^ for the first 4 days for all monkeys). Furthermore, not all monkeys succeeded in generalizing this learned rule to novel texform images (Fig. 3D; Monkeys Ju and Ol: *p <* 2.1 *×* 10^−5^, Monkey El: *p* = 0.078, BF_01_ = 1.9, binomial test compared against chance level). Therefore, the monkeys’ classification depended strongly on higher-level features than those preserved in texform.

As yet another test of whether monkeys learned specific stimuli instead of learning a common rule, we devised two versions of control tasks in which we associated random object images with targets (Fig. 3E, F). In one version, all new images (*n* = 96) were randomly assigned to two categories, thus monkeys had to remember stimulus-target associations individually (Fig. 3E). To our surprise, the monkeys still gradually learned these many associations over a few days, but their learning speed was much slower than in the main tasks (Fig. 3G; *p <* 10^−10^ for all days, aggregated across monkeys, binomial test). In the second version, we randomly assigned object categories to two targets, but different images within the same concrete category (e.g., two images of bottle) were always associated with the same target (Fig. 3F). Nonetheless, the learning speed was again slower than the main tasks (Fig. 3G), and importantly, the monkeys performed poorly in generalizing these category associations to new images (Fig. 3H; *p <* 0.016 for all three monkeys, binomial test). Thus, the monkeys did not categorize new images purely on the basis of similarity to individual images they had learned.

**Figure 3:**
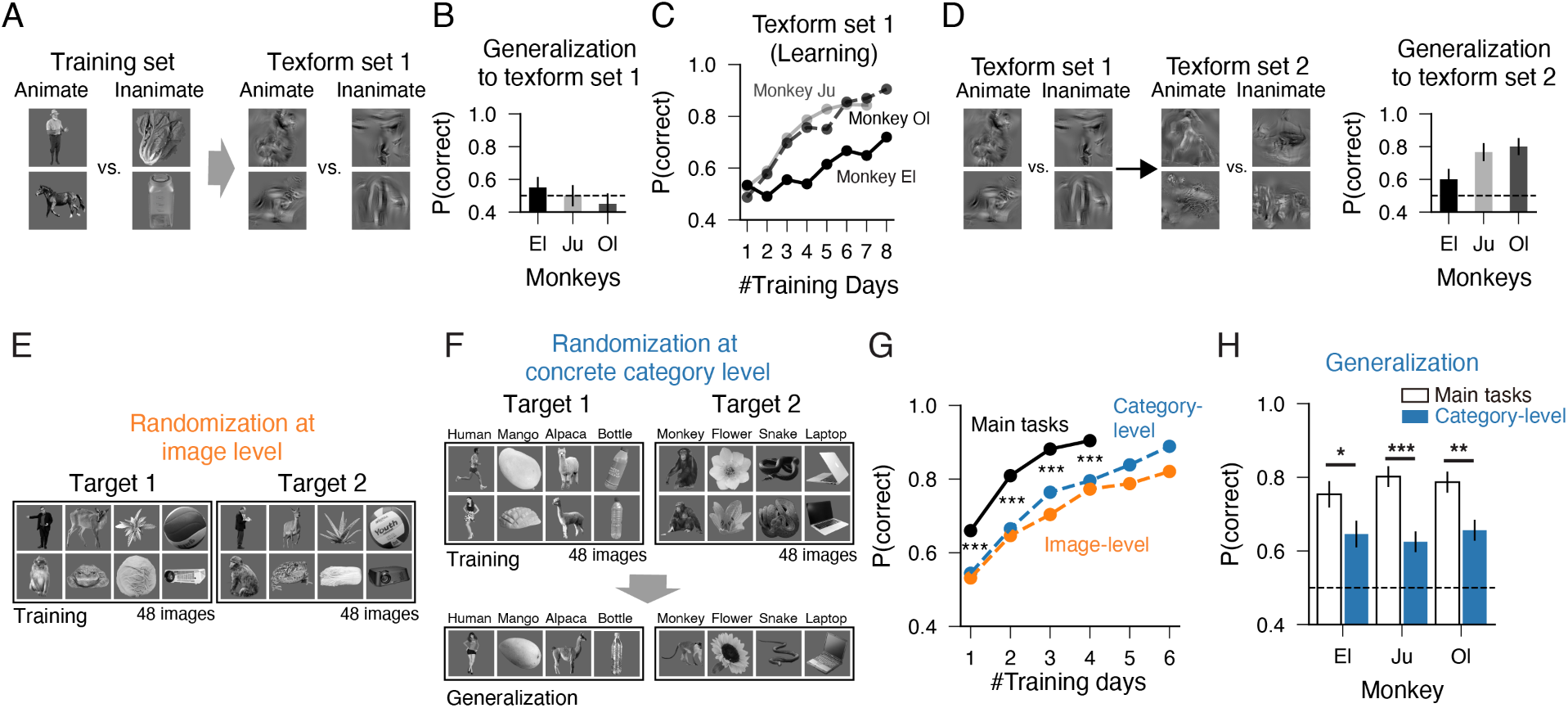
Lack of high-level image features or absence of a common conceptual rule results in poorer performance. (**A**) We tested whether monkeys generalize the animate vs. inanimate rule to “texform” stimuli, which preserve mid-level textural visual features informative for animacy recognition (Long et al. 2018) but lack higher-level image structure. (**B**) Monkeys’ performance on the texform stimuli was at chance level. (**C**) We then attempted to train the monkeys to discriminate the texform version of animate vs. inanimate (Set 1). Although the monkeys gradually learned it, their learning speed was slower than that in the main tasks. See Supplementary Fig. 3 for the learning time courses as a function of trial count. (**D**) Monkeys Ju and Ol generalized the learned rule to novel texform images, whereas monkey El’s performance stayed near chance level. (**E**) A random association task, in which two targets were paired with randomly selected images (48 images per target) with no common rule, requiring monkeys to memorize the associations for each individual image. (**F**) Another randomization task, in which two targets were paired with concrete object concepts (e.g., apple, horse, bin; 16 concrete categories for each target). In the generalization test, different images of the same concrete categories were shown. (**G**) Although the monkeys could gradually learn both control tasks, performance was consistently lower than on the main tasks (black line; the average of the six tasks shown in Fig. 1). (**H**) Their generalization performance for the concrete category randomized task (F) was also lower than that of the six main tasks (p < 0.016 for all three monkeys, binomial test). Error bars are S.E.M. across images. Asterisks indicate statistical significance: ^∗^p < 0.05, ^∗∗^p < 0.01, ^∗∗∗^p < 0.001.

### Testing on more diverse categorization rules

Thus far, monkeys have generalized various natural categorization rules we have tested (Figs. 1, 2), but are there any rules they cannot generalize? To explore this, we further created five additional categorization tasks that challenge them (Fig. 4A). As often investigated in the vision science literature, we first tested outdoor vs. indoor scenes (Zhou et al. 2017) and big vs. small object categorization (Konkle and Oliva 2012; Long et al. 2016). Big vs. small objects were determined on the basis of physical object size (objects bigger than a chair were classified as “big”; Konkle and Oliva 2012) and had no correlation with image size. We had two versions of big vs. small, one using the set from Konkle and Oliva (2012) and one using images from the THINGS database. We then created two more tasks that would rely heavily on human knowledge: “fire-related” vs. “water-related” objects, such as lighters, fire extinguishers, faucets, and bathtubs, and “Western culture” vs. “Eastern culture” objects, such as mooncakes, Chinese knots, crowns, and Easter eggs. For each task, we first trained the monkeys on training images for 4-6 days and then assessed their generalization performance on test images as in Fig. 1B.

The monkeys successfully generalized the outdoor vs. indoor rule to new stimuli (Fig. 4B; *p <* 4.3 *×* 10^−7^ for all monkeys, binomial test) and also performed well on the two versions of the big vs. small task (*p <* 2.4 *×* 10^−6^ for all monkeys in both tasks, binomial test). In contrast, their performance was substantially lower for the fire- vs. water-related task (46-64% correct) and their performances were at chance level for the Western vs. Eastern object task (*p* = 0.38 *−* 0.86, BF_01_ = 4.9 *−* 7.4 across monkeys, binomial test). By contrast, we found that human participants generalized these rules almost perfectly, even for the latter two rules using the same training and generalization images (Fig. 4B, black line; see Methods), demonstrating a clear human advantage on these tasks.

**Figure 4:**
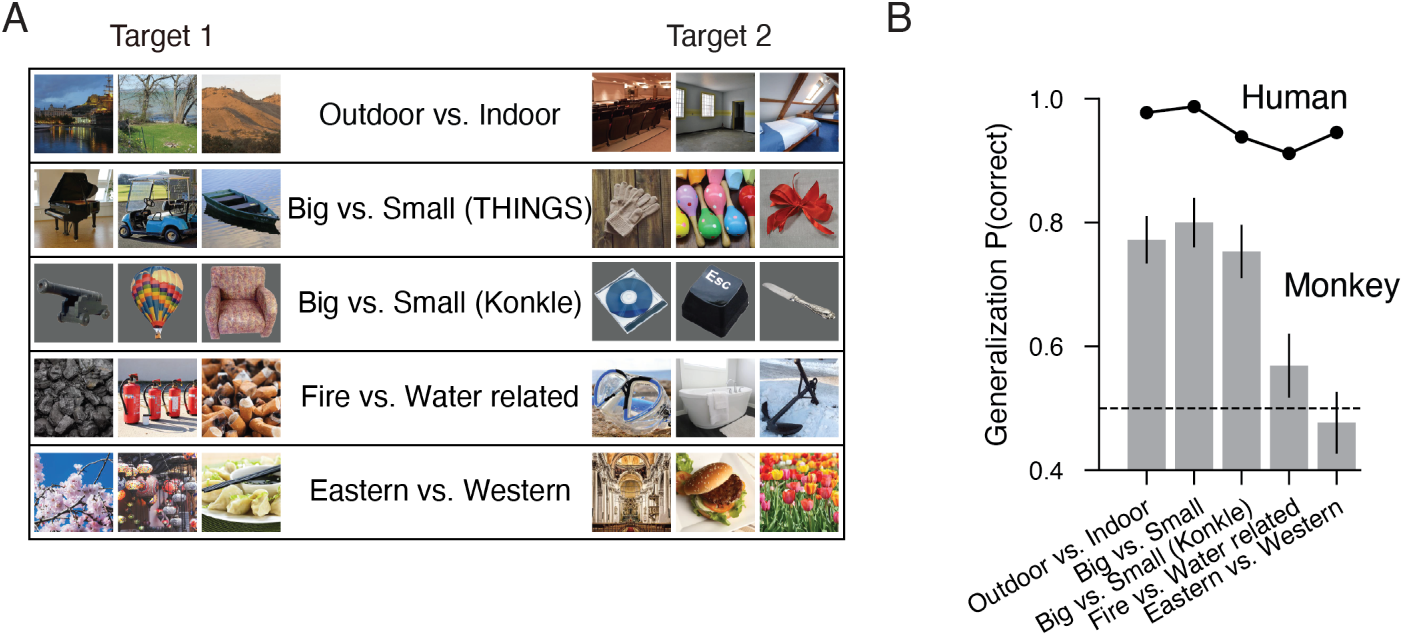
Testing on more diverse categorization rules. (**A**) To determine the limit of monkeys’ ability to classify concepts, we tested additional concept tasks. As with the other tasks in Fig. 1, we trained the three monkeys with 90-120 images and assessed their generalization performance with 90-120 new images. (**B**) The monkeys (n = 3) performed well on tasks such as big vs. small objects and outdoor vs. indoor scenes, but they failed on some abstract tasks, such as fire- vs. water-related objects and Western vs. Eastern objects. Human participants (n = 9) performed near perfectly on all tasks.

### Features informative for solving the tasks: comparison with vision models

What visual features allow such category generalization and to what extent can deep network models (DNNs) perform the same tasks? To test this, we created a range of vision models from low-level image features to state-of-the-art DNNs (Fig. 5). Low- and mid-level image models included V1-like filter outputs (Pinto et al. 2008), luminance- and color-based statistics, spatial-frequency summary statistics, and texture statistics (Portilla and Simoncelli 2000). DNNs included AlexNet (Krizhevsky et al. 2012), VGG16 (Simonyan and Zisserman 2014), ResNet-50 (He et al. 2015), and Vision Transformer (ViT-B/32) (Dosovitskiy et al. 2020), which were pretrained on ImageNet. We also tested a self-supervised vision model, DINO (Caron et al. 2021), and two recent large-scale models (CLIP (Radford et al. 2021) and SigLIP2 (Tschannen et al. 2025)) that were trained to match visual and text inputs, thus providing additional language supervision. For all the models, we first reduced their output dimensions using principal component analysis (PCA), constructed a linear classifier for either the true categories or the monkeys’ choices for the training set, and then tested their performance on the generalization set. The results did not depend on specific details of the dimension-reduction and regression methods (Supplementary Fig. 4).

For the three main THINGS tasks (Fig. 2), we found that all DNNs performed the tasks and fit monkey behavior well, whereas low-level vision models did not (Fig. 5A, B). The classification performance of the low- and mid-level models was typically *∼*60%, and they were all significantly lower than the monkey performance (*p <* 10^−10^ for all comparisons between models and average monkey performance, binomial test, Bonferroni-corrected for multiple comparisons).

Correspondingly, even when these models were fit directly to monkey behavior, the rates of choice agreement between the models and the monkeys were low (*∼*60% in many cases). By contrast, all DNNs performed our tasks with high accuracy (*>*90%) and fit the monkey behavior better than the low- and mid-level visual models (*p <* 10^−10^ for all comparisons between models; binomial test, Bonferroni-corrected for multiple comparisons).

When we tested DNNs on more categorization tasks, we found interesting similarities and dissimilarities in performance between monkeys and different DNNs (Fig. 5C, Supplementary Fig. 5). While all DNNs performed well on most tasks, conventional DNNs trained without language information (e.g., AlexNet, ResNet-50) suffered on abstract tasks such as the fire- vs. water-related task (53-75% correct across models) and the Western vs. Eastern task (46-60% correct across models), mirroring the monkey behavior. By contrast, the performance of a language-informed model (e.g., SigLIP2) was better (*p <* 6.9 *×* 10^−6^, binomial test compared against chance level).

For the control conditions such as silhouette, texform, and rule randomization, we also found that visual DNNs performed qualitatively similarly to monkey behavior (Supplementary Fig. 6).

We further leveraged previous large-scale neural recordings from monkey visual areas along the ventral pathway (V1, V4, and IT cortex) using the same THINGS image set (Papale et al. 2025) and tested neural classification performance on our tasks (Fig. 5D). We first trained a linear classifier on the training set for each task and then tested performance on the generalization set, following a procedure of the monkey experiments. As expected, performance was higher for later stages of the ventral pathway (Fig. 5D) and improved with the number of units included. However, even with hundreds of units, IT cortex performance did not reach monkey-level performance on some tasks. To test whether this reflects a limitation in IT-level visual coding, we used an alternative classification procedure: 5-fold cross-validation within the generalization images, which allowed many more images to be used for training the classifiers. This procedure yielded IT performance comparable to that of the monkeys (Fig. 5E). Thus, IT cortex contains sufficient information to solve these tasks, and the monkeys’ higher performance with limited training samples (Fig. 5D) may be due to a better learning strategy or prior experience.

**Figure 5:**
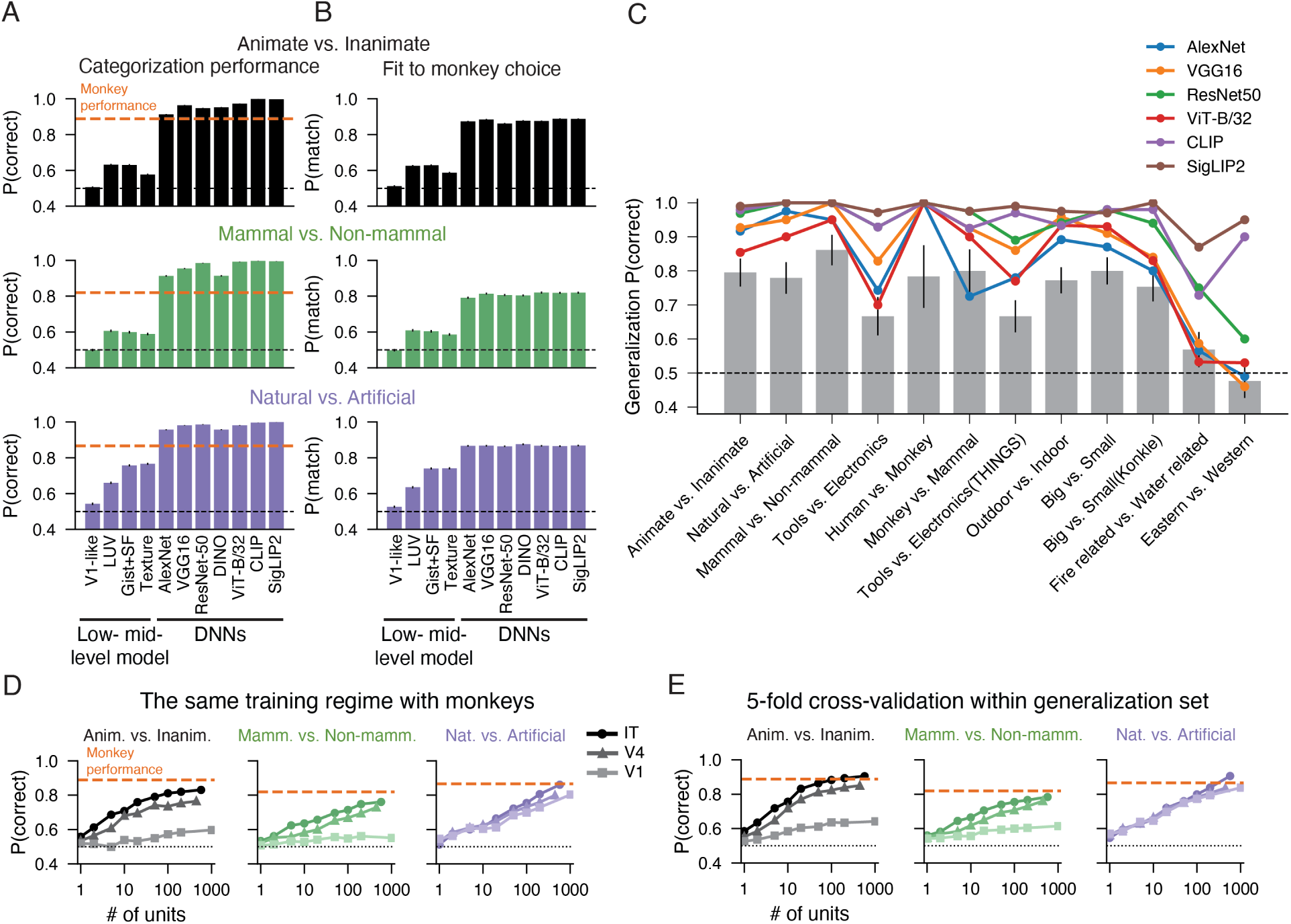
The categorization rules the monkeys learned could also be classified by visual DNNs and IT neural responses but not by low- or mid-level vision models. (**A**) Category classification accuracy of various vision models, including deep neural networks (DNNs) for the three THINGS tasks in Fig. 2. We constructed a linear classifier using the output of each model based on the training images and plotted their performance on the generalization images we used for the monkeys. The orange dashed lines represent the average classification performance of the three monkeys. (**B**) Accuracy of predicting monkey choices. A linear classifier was trained to classify monkey choices instead. P (match) indicates the probability that the monkey and model choices matched. DNNs outperformed lower-level vision models. (**C**) Full comparison of monkey and DNN generalization performance across a test battery. Both monkeys and visual DNNs performed well on most tasks other than abstract categorization rules (e.g., Western vs. Eastern), whereas language-informed DNNs (e.g., SigLIP2) could also perform these tasks. (**D**) Accuracy of category classification based on neural responses to THINGS images in ventral pathway visual areas, calculated from data collected by Papale et al. (2025). Linear decoders were trained on the training set and tested on the generalization test, following the monkey experiment procedure. (**E**) IT classification performance reached monkey-level performance when decoders were trained with 5-fold cross validation within the generalization images, which provided more training samples.

### Image-level monkey performance was correlated with human performance and typicality

We next sought to compare monkey and human behavior in the same tasks shown in Fig. 1 to determine whether they share similar categorization patterns at the level of individual images. When human participants performed the same tasks, they understood the rules within tens of trials and thereafter showed near-perfect performance across all tasks (Fig. 6A), making it impractical to compare patterns of correct rates. However, they still showed different reaction times (RTs) across images (Fig. 6B, *F* (95, 570) = 2.88, *p <* 10^−10^ for animate vs. inanimate, *F >* 2.14, *p <* 4*×*10^−6^ for the other tasks, repeated-measures one-way ANOVA). Thus, we expected that difficult stimuli would cause longer RTs in humans and lower choice accuracy (and longer RTs) in monkeys (Fig. 1E-F).

We therefore constructed a metric of stimulus difficulty that incorporates both choice accuracy and RTs. We used the drift-diffusion model (Fig. 6C), which has been widely used to explain choices and RTs in various experimental settings (Carlson et al. 2014; Heidari-Gorji et al. 2021; Luo et al. 2025; Mack and Palmeri 2011). The model assumes a one-dimensional stochastic internal state that meanders between two decision bounds corresponding to two choice options until it reaches one of them (Fig. 6C). An easy stimulus causes a large drift (higher sensitivity) toward the correct bound, leading to higher accuracy and shorter RTs. The slope of the drift is termed “category sensitivity” here, whose absolute value reflects how easily an image can be categorized correctly. Note that we did not assume that an evidence accumulation process underlies our task (Stine et al. 2020). Rather, we used the model as a means of deriving a single metric of stimulus difficulty (the absolute value of category sensitivity) for each image from the choices and RTs. Critically, the model fits both human and monkey data well (Fig. 6D, Supplementary Fig. 7; *R* = 0.99 *−* 1.0 for choices and *R* = 0.81 *−* 0.99 for RTs across monkeys, humans, and tasks), demonstrating its utility.

For all tasks, the absolute value of the category sensitivity of humans and monkeys across images had moderate positive correlations (Fig. 6E, F). Humans obviously had higher category sensitivity for all images, as expected from their near-perfect performance. Nonetheless, there was a tendency for images with higher category sensitivity for humans to also be higher for monkeys (Fig. 6E). The correlations of sensitivities between humans and monkeys were significantly positive for all tasks (Fig. 6F; *R* = 0.30 *−* 0.43; *p* = 7.4 *×* 10^−5^ *−* 6.0 *×* 10^−3^). The positive correlation in performance between the two species also extended to images outside the range of natural photographs (Fig. 6G, H), such as the cartoon, silhouette, and outline images used in the animate vs. inanimate task (part of the results shown in Fig. 2D). The monkeys performed better on cartoon and silhouette images than on outline images, consistent with the lower human category sensitivity observed for outline images (Fig. 6G).

**Figure 6:**
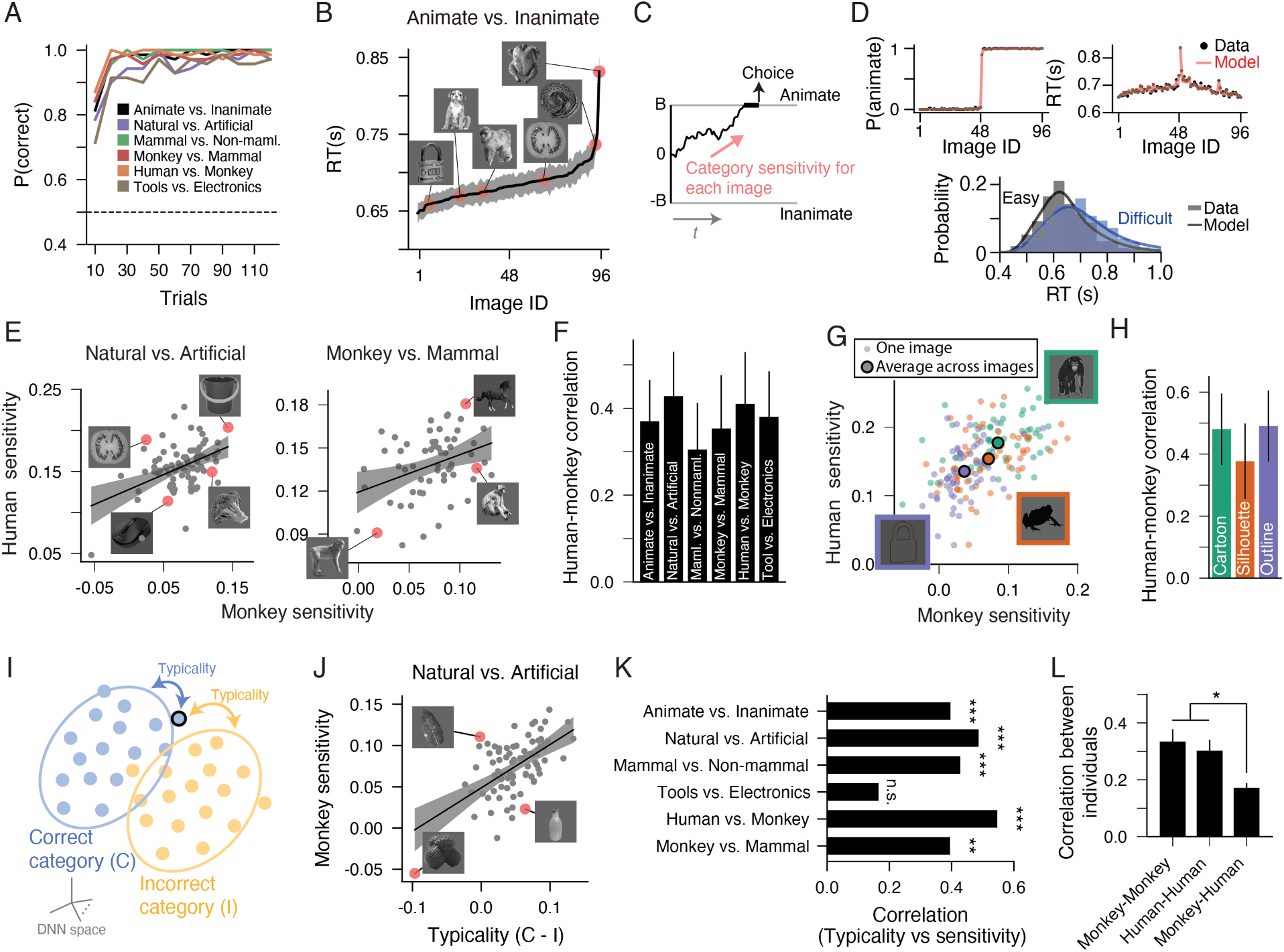
Human behavioral responses were correlated with monkey performance. (**A**) Humans quickly learned the tasks in Fig. 1 within 20-30 trials. The plots are the average of seven participants per task. (**B**) Reaction times differed across images (averaged across participants). (**C**) We fitted the drift-diffusion model (DDM) to the choices and RTs. The model makes a decision based on a stochastic state biased by the category sensitivity of each image. (**D**) DDM accurately fits choices and RTs. The plots are from the animate vs. inanimate task. Image IDs were sorted by category sensitivity. RT distributions were generated using the trials aggregated across the top (easy) and bottom (difficult) 10 images. See Supplementary Fig. 7 for all tasks. (**E**) Example comparison of category sensitivity between humans and monkeys. Each dot represents an image, whose absolute category sensitivity was averaged across participants. The line is a linear regression, and shading indicates S.E.M. of the prediction line. (**F**) Positive correlations between humans and monkeys. Error bars indicate S.E.M. across images. (**G**) The category sensitivity for control images —cartoon, silhouette, and outline— was in the same order between monkeys and humans. (**H**) Positive correlations were also evident within each control image set. (**I**) To test whether the monkeys made more errors for atypical images, we calculated the typicality of each image between two categories in each task based on DNN outputs (Kramer et al. 2023). (**J**) Example comparison of monkey category sensitivity (average of all monkeys) and image typicality (calculated as the difference in typicality for the correct and incorrect category) in one task. (**K**) Monkey category sensitivity was significantly correlated with image typicality in most tasks. ^∗∗^p < 0.01, ^∗∗∗^p < 0.001. (**L**) Between-subject correlations of category sensitivity were higher within each species than across species. ^∗^p < 0.05.

These results naturally led us to hypothesize that the monkeys perform poorly on less typical exemplars within each classification rule. To quantitatively confirm this, we followed Kramer et al. (2023) to compute the typicality of each image using the outputs of DNNs (Fig. 6I). We calculated pairwise correlations of DNN outputs across all images in a task and, for each image, averaged its correlation with all other images in the same (correct) and different (incorrect) categories, and took their difference (e.g., a dog image was compared with all other animate and inanimate images). There were significant correlations between this image typicality and the monkeys’ categorization sensitivity in all tasks (Fig. 6J, K; *R >* 0.39, *p <* 0.002; calculated using AlexNet, and other visual DNNs yielded similar results) except for the tools vs. electronics task (*R* = 0.16, *p* = 0.15). The effect of typicality was also seen in generalization performance on THINGS images (Supplementary Fig. 8), supporting the idea that image typicality strongly influenced the monkeys’ categorization behavior.

At the same time, we also found evidence of differences between human and monkey judgment.

When we compared sensitivity across individual subjects, rather than averaging across subjects, we found correlations within each species (Fig. 6L; humans, *R* = 0.30, *p* = 7.8 *×* 10^−4^; monkeys, *R* = 0.33, *p* = 8.4 *×* 10^−4^), but the correlations between individuals of different species were significantly lower (*R* = 0.17, both *p <* 0.018, two-tailed *t*-test).

### Triangular comparison across monkeys, humans, and artificial networks

As a collective summary of the results, we combined generalization performance across all tasks to visualize the overall similarity of behavioral responses among monkeys, humans, and vision models (Fig. 7). For monkeys and humans, the average correct rate was calculated for each generalization image across subjects and then concatenated across tasks. For the models, we used the likelihood of correct classification for each image based on a logistic classifier trained on the training images. A matrix of dissimilarities among humans, monkeys, and models based on these values (Fig. 7A) revealed that monkey choices were closer to DNNs trained without language input (e.g., AlexNet; Fig. 7B bottom), whereas human choices were closer to the language-informed DNNs (CLIP and SigLIP2; Fig. 7B top). Consistent with this, t-SNE visualization revealed a spectrum from low-level visual models (e.g., the V1 model) to language-informed DNNs, with the monkey data located in the middle, proximal to visual DNNs (Fig. 7C).

**Figure 7:**
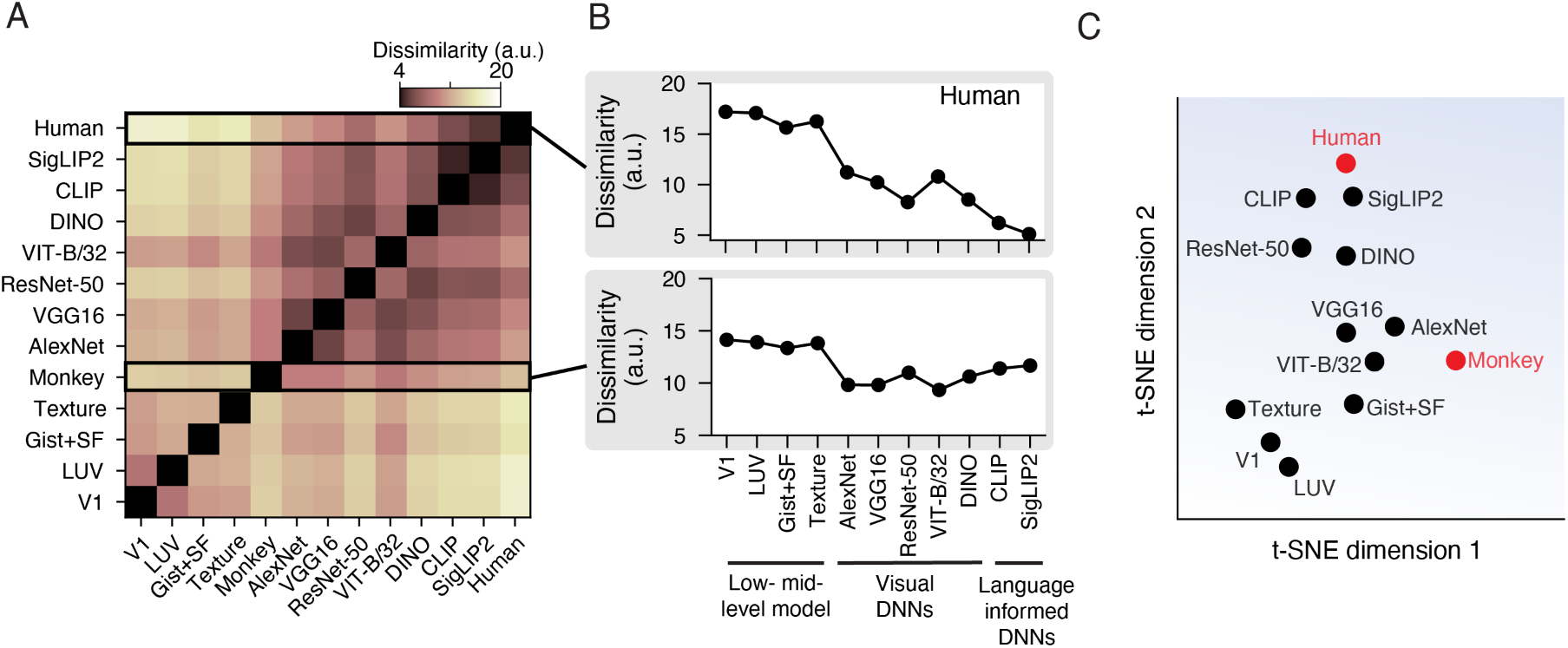
Triangular comparison across monkeys, humans, and artificial networks. (**A**) Finally, we concatenated the generalization performances of images across all tasks to generate the matrix of dissimilarity in behavioral performance across humans, monkeys, and models. (**B**) A closer look at the dissimilarity revealed that the monkey behavior was most similar to visual DNNs without language input, whereas the human behavior was most similar to language-informed DNNs. (**C**) t-SNE visualization of the dissimilarity matrix showed a spectrum from low-level visual models to language-informed DNNs, with the monkey behavior placed in the middle.

## Discussion

Macaque monkeys serve as a primary model for investigating the human visual system, allowing characterization of neural mechanisms at the cellular level and providing insight into the evolution of object vision (Orban et al. 2004; Yamins and DiCarlo 2016), but it remains poorly understood whether monkeys can classify objects according to rules at various levels of abstraction. We developed a task design that enabled us to test monkeys using a battery of image-classification challenges to quantify their capacities and limitations. Monkeys generalized a surprising variety of categorization rules to novel images, and their behavior could not be explained by learning individual objects (Figs. 2, 3) or by classification based on low-level image features (Figs. 2D, 5A). Instead, monkey behavior was best fit by visual DNN outputs (Fig. 5A, B) and was also partially correlated with human behavioral responses (Fig. 6). At the same time, monkeys failed to learn some abstract rules (Fig. 4B). Overall, whereas human behavior was proximal to language-informed DNNs, monkey behavior was closer to DNNs without language input (Fig. 7). The present study represents an unprecedentedly systematic characterization of animals’ object-classification performance.

The widespread use of macaque monkeys in ventral pathway research was guided by the premise that they recognize objects just as humans do, but there is not even consensus on how to measure their object-recognition behavior. One approach is to use tasks that require monkeys to report the similarity of object images, such as match-to-sample (Rajalingham et al. 2015, 2018) and odd-one-out (Cherian et al. 2025; Hebart et al. 2020). Since these tasks do not require training of individual categories, the similarity can be measured across many combinations of object categories to generate confusion matrices. The downside, however, is that the methods lack explicit control over which properties of object images the animals rely on to report similarity. Indeed, confusion matrices measured in match-to-sample (Rajalingham et al. 2015) do not appear to be strongly influenced by higher-level categories such as animacy (but see Cherian et al. 2025 for the odd-one-out case). The second approach is to explicitly train animals to classify two categories (e.g., dog vs. cat) and measure their generalization performance, but in this case, each task must be trained individually. Consequently, previous studies performed only one or two tasks chosen according to their experimental purposes (Eldridge et al. 2018; Fabre-Thorpe et al. 1998; Minamimoto et al. 2010; Vogels 1999). Our innovation is that by leveraging a rapid training paradigm, we scaled up the second approach into a large test battery. While we focused on various natural-object concepts used in vision research, the rules do not have to be confined to verbally describable categories or natural-object images.

It was a surprise to us that the monkeys learned each task rule quickly, although the tasks required them to discover a common classification rule from many object images with different appearances, which contrasts with conventional visuomotor association tasks that use only a few images (Cromer et al. 2011). We also observed rather surprising consistency in the monkeys’ learning curves across a variety of classification tasks (Fig. 1H). Many studies posited that basic-level categories are most accessible due to the family resemblance of image features within a category (Grill-Spector and Kanwisher 2005; Rosch et al. 1976), whereas other studies have reported that superordinate categories are processed faster in the human brain (Clarke and Tyler 2015; Macé et al. 2009). But these differences in speed might simply depend on the amount of evidence needed to make a decision (Mack and Palmeri 2011), and the underlying decision-making mechanisms might not qualitatively differ across levels of object classification.

Our paradigm does not answer whether the monkeys were truly aware of object concepts such as “animacy” and “natural”, but the results still provide insights into the mechanisms that underlie the formation and representation of such concepts in the brain. Recent works have proposed that conceptual representations are similar between humans and DNNs, based on alignment in their internal representations (Du et al. 2025; Muttenthaler et al. 2025; Vong et al. 2024). However, to our knowledge, there is no established definition of what “object concepts” entail. One may regard multimodal associations (Quian Quiroga et al. 2005) or arbitrary sensory associations (Loggia et al. 2025), which cannot be explained by mere perceptual similarity, as key traits of object concepts. Earlier psychological studies also emphasize the importance of knowledge structure, such as recognizing the functions of objects to form concepts (Murphy 2004). However, many of the object concepts we used here were decodable from natural images through visual DNNs and could be classified by monkeys. These results rather support the view that many object concepts are already structurally embedded in retinal images to some extent (Vong et al. 2024) and that the primate brain can readily learn to extract them even without non-visual knowledge. While there should be additional contributions from knowledge and multimodal associations, it is conceivable that such embedding of concepts in images can substantially facilitate the formation of object concepts in the human brain.

## Acknowledgment

We thank Takafumi Minamimoto and Cheng Xue for comments on earlier versions of the manuscript. We thank Roozbeh Kiani for providing natural object images and also thank the Japan Monkey Centre and the Center for the Evolutionary Origins of Human Behavior, Kyoto University, for providing monkey images used in the experiments. This work was supported by Strategic Priority Research Program of the Chinese Academy of Sciences (XDB1010202), the National Science and Technology Innovation 2030 Major Program (Grant No. 2021ZD0203703), National Natural Science Foundation of China (Grant No. 32371077 and No. W2432019), and Shanghai Municipal Science and Technology Major Project (Grant No. 2019SHZDZX02).

## Data and code availability

Data and code used in this study are available at https://github.com/okazawagouki/Zhang_Okazawa_ObjectClassification

## Methods

### Subjects and apparatus

#### Monkey experiments

We collected behavioral data from three adult rhesus monkeys (*Macaca mulatta*; monkey Ju, female, age 13; monkey El, female, age 12; monkey Ol, male, age 11). These monkeys were raised in a typical laboratory environment and had no prior experience performing relevant behavioral tasks. The dataset collected in this study consisted of 26 tasks including subtasks, amounting to *∼*315K trials from the three monkeys. Experimental procedures conformed to the National Institutes of Health *Guide for the Care and Use of Laboratory Animals* and were approved by the Animal Care Committee at Center for Excellence in Brain Science and Intelligence Technology (Institute of Neuroscience), Chinese Academy of Sciences.

Experiments were conducted using a custom-built touchscreen device attached to each monkey’s cage (Supplementary Fig. 1A), similar to those developed by other laboratories (Butler and Kennerley 2019; Ramezanpour et al. 2024). The device box consisted of a commercial touchscreen (AOSIMAN, ASM-125UCT, 12.5 inches; 60 Hz refresh rate; 3840 × 2160 pixel screen resolution), a reward juice delivery system with a TTL-triggered water pump, a monitoring camera, and a control computer. The monkey reached the touchscreen through a hole in the front panel attached to the cage and viewed the screen through another hole, which opened at approximately the level of the monkey’s head. Below the hole was a reward juice outlet in a “cone”-shaped socket of a size suitable to cover the monkey’s mouth (Kawaguchi et al. 2019). The monkeys performed the tasks while their mouth was in this socket and thus had a stable head position. Under these conditions, the viewing distance to the monitor was approximately 15-18 cm. The task was controlled by a custom-made MATLAB (MathWorks, MA, USA) program with Psychtoolbox.

#### Human experiments

We recruited in total 33 human participants (20-40 years old; 10 males and 23 females, students or employees at the Chinese Academy of Sciences). Our participant sampling did not consider sex, as it was considered unlikely to influence the results of this study. They were assigned to perform different tasks, as explained below. All participants had normal or corrected-to-normal vision and were naïve to the purpose of the experiments. Written informed consent was obtained from the participants. All experimental procedures were approved by the Institutional Review Board of the Center for Excellence in Brain Science and Intelligence Technology (Institute of Neuroscience), Chinese Academy of Sciences.

During the experiments, the participants were seated in a height-adjustable chair, with their chin and forehead supported by a chinrest. The chinrest ensured a stable viewing distance (37 cm) to a touchscreen monitor (LENOVO, T24t-20, 23.8 inches; 60 Hz refresh rate; 1920 x 1080 pixel screen resolution). The viewing distances were different from those used in monkey experiments, but we maintained similar visual angles of the object images (see below). The other task parameters were also kept identical to those of the monkey experiments.

### Object drag task

We developed a binary object categorization task (“object drag task”) for the touchscreen device (Fig. 1A). During the task, the participants viewed an object stimulus and moved it to one of two target boxes by swiping it with their fingers. The two target boxes were associated with distinct object concepts (e.g., animate vs. inanimate objects).

Each trial started when the participants touched a red fixation dot. Following this, a stimulus (10 *−* 11 deg) appeared together with two gray square target boxes. The positions of the stimulus and target boxes were adjusted for each participant so that they could move a stimulus smoothly to both targets with similar amounts of time. In the monkey experiments, the distance between the stimulus and target boxes was 18 *−* 24 deg, and the distance between the two target boxes was 24 *−* 37 deg. Following the stimulus presentation, the participants were required to touch the stimulus within 1 s. After the touch, the participants were required to drag the stimulus to one of the two target boxes within 5 s. In the case of timeout, the trial was aborted (1.3 % of trials for monkeys). When the image reached one of the two boxes, the image and the target boxes disappeared, and instead, the same image appeared at the correct target location, providing visual feedback to the participants. At the same time, an auditory tone indicated correct and incorrect responses. The monkeys received a juice reward for a correct response and a short timeout (1-2 s) for an incorrect response.

A rule for a given task (e.g., animate vs. inanimate) did not change within a daily experimental session for the monkeys (see below for rules). The same rule usually lasted for one to two weeks until the participants learned the task and completed a generalization test (Fig. 1B). At the start of a new rule, they were not informed of the rule change, but the new task involved different stimuli, which may have reduced the interference from the previous task and facilitated the learning of the new rule.

### Image sets

#### Background-cropped image sets

In the first series of our experiments (Fig. 1), we used grayscale object images with the background removed (examples shown in Fig. 1B, G). Some of the images (5%) were obtained from an existing database (Kiani et al. 2007), whereas the remaining images were obtained by manually removing background from natural photographs of objects obtained through search engines such as Bing, Pixabay, and Google Images. We also obtained photographs of monkeys from the Japan Monkey Centre and the Center for the Evolutionary Origins of Human Behavior, Kyoto University. Images were converted to grayscale because the distribution of colors was highly skewed in some objects (e.g., monkey images are always brownish). Because some original images do not allow redistribution, we replaced them with similar images with the Creative Commons Zero (CC0) license or other royalty-free licenses when shown in the figures. The sources of the human images shown in Figs. 1, 3 are PurePNG (https://purepng.com/; authors: LPuo, hadiaaltaf, Momo, FMPure).

When collecting images for these sets, we focused on eight object groups: humans, monkeys, non-primate mammals, non-mammals (reptiles/amphibians), food, plants, tools/everyday objects, and electronics. Although these eight groups span different levels of category hierarchy (e.g., “monkey” can be considered a basic category, whereas “food” is a superordinate category), we considered that these groups covered a range of object categories ideal for behavioral experiments with monkeys.

Using these eight object groups, we created six tasks (Fig. 1B, G): animate vs. inanimate, natural vs. artificial, non-primate mammals vs. non-mammals (reptile/amphibian), human vs. monkey, monkey vs. non-primate mammal, tool/everyday object vs. electronics. In each task, 60-96 images were used to train the monkeys, and 20-96 images were used to test their performance on novel stimuli after learning (generalization test; Fig. 1B). Training images could have a small number of overlaps across tasks, whereas the generalization images were never shown to the monkeys in any previous tasks.

#### Large-scale THINGS image sets

To complement the experiments using the controlled but small stimulus sets above, we created large-scale image sets based on the THINGS database (Hebart et al. 2019, 2020; Stoinski et al. 2024). The THINGS database consists of 1,854 concrete object concepts (such as dog, spider, lemon, and table), each of which contains at least 12 natural photographs.

For the purpose of our experiments, we selected a subset of object categories from the THINGS database and created three tasks (Fig. 2A): animate vs. inanimate (373 object categories, 3,683 images in total), natural vs. artificial objects (230 object categories, 1,915 images in total), and mammal vs. non-mammal animals (143 object categories, 1,668 images in total). Among the images, we selected 100 images (50 images per target; no overlap in object categories) for training the monkeys on each task. During the generalization test, we showed the remaining images only once each. Due to copyright issues, the images shown in the figures are replaced with similar images with the CC0 license from the THINGSplus database (Stoinski et al. 2024) or images with the Pixabay Content license from Pixabay.

To inspect the patterns of monkey behavior, we further classified these objects into high-level object categories such as “bird”, “plant”, and “fruit” (Fig. 2C). These high-level categories were mostly taken from the THINGS database (Hebart et al. 2019), which uses human curation to determine them. In addition, we included several high-level categories that we defined ourselves (mammals, non-mammals, natural, artificial, reptiles/amphibians, fish, and Mollusks/Crustaceans) as they could be informative for assessing monkey performance.

#### Modified control image sets

After performing the animate vs. inanimate task using the grayscale images (Fig. 1), we tested whether the monkeys could classify images outside the range of natural photographs (Figs. 2D, 6G, H). We created three control image sets. First, an “outline” image set was created by extracting the outlines of the object images using the *bwboundaries* function in MATLAB. The contour was colored black, while its interior was filled with the background color of the task screen (gray). Second, we created a “silhouette” image set, where the inside of a contour was also filled with black. Third, we created a “cartoon” image set by selecting cartoon images of objects from a publicly available database (the “Multipic” database; Duñabeitia et al. 2018). In all the sets, we selected source images that were not shown during the training to ensure that the monkeys could not solve the task by remembering specific object images. Each set contained 60 images.

We also presented two control image sets after performing the categorization of large-scale THINGS images. First, during the natural vs. artificial task, we showed THINGS images converted into grayscale to confirm that the monkeys did not simply rely on color. Second, during the mammal vs. non-mammal task, we presented THINGS images whose texture was distorted by using *crystallization* function (cell size, 10) in Photoshop 2024 (Adobe Inc., San Jose, CA, USA). We had 400 images for each set, selected from the generalization image set. These images were shown once during the generalization test, but we consider it unlikely that the monkeys remembered specific images to solve these control tasks because they were shown only once and mixed with many other images.

#### Image sets for more conceptual tasks

To further challenge the monkeys’ ability to learn abstract rules, we created additional conceptual tasks (Fig. 4). Each task used 90-120 training images and 90-120 generalization images.

##### Outdoor vs. indoor scenes

We selected scene images from Zhou et al. (2017). The indoor scenes included pictures of a variety of buildings, including small scenes such as bedrooms and kitchens and also spacious scenes such as stadiums and concert halls. The outdoor scenes included both natural scenes, such as forests, and urban scenes, such as buildings. Because the sky and natural objects (e.g., a tree) could be strong cues indicating outdoors, we preferentially selected images that did not contain them (e.g., the front face of a building). Nevertheless, outdoor scenes tend to have certain colors (greenish). Therefore, we created a grayscale version of the generalization images as a control and confirmed that the monkeys also performed well without color (73 % correct).

##### Big vs. small objects

In this task, the monkeys had to classify objects based on their physical size. Objects that are typically bigger or smaller than the size of a chair were classified as big or small, respectively (Konkle and Oliva 2012; Long et al. 2016). We created two sets of stimuli. The first set used images from the THINGS database, which had a natural background. Small objects included many household items such as spoons and nails, and big objects included furniture, vehicles, and others. We ensured that pictures of most big objects were taken indoors so that this categorization could not be confused with outdoor vs. indoor. The second set used images from the big and small images created by Konkle and Oliva (2012), which had no background. A comparison of these two sets allowed us to confirm that the monkey performance did not depend on a specific choice of images.

##### Fire- vs. water-related objects

To create abstract rules that are unlikely to be correlated with any visual features, we selected object images that are semantically associated with fire and water from the THINGS database. The fire-related category included objects such as stoves, cigarettes, coal, fire trucks, and radiators. The water-related category included objects such as hoses, bathtubs, paddles, and squirt guns. The images of fire or water itself were avoided as they have distinctive features.

##### Eastern- vs. Western-culture objects

As another abstract categorization task, we created an image set of objects associated with Eastern and Western cultures. We collected the images mainly by using Internet search engines, because a sufficient number of images could not be found in the THINGS database. The Eastern category included objects such as Chinese lanterns, dumplings, old pavilions, bamboo, and chopsticks. The Western category included objects such as crowns, donuts, the Eiffel Tower, tulips, and violins.

##### Tools vs. electronics (THINGS)

To test whether the monkeys performed similarly with natural photographs as with the background-cropped grayscale image set, we also created a tools vs. electronics task using THINGS images. Each category included 50 training images and 50 generalization images. The tools category included objects such as hammers, clippers, and saws. The electronics category included objects such as cellphones, headsets, and radios.

#### Texform images

Texforms are synthetic images generated by matching the mid-level texture features of natural object images (Long et al. 2016, 2017, 2018). These features consist of the magnitudes and correlation structures of V1-like oriented bandpass filter outputs and some other image statistics at each local image region (Freeman et al. 2013; Portilla and Simoncelli 2000). Humans can reliably classify the animacy of original objects from texform images (Long et al. 2017); thus, we tested whether monkeys could generalize the animate vs. inanimate task to texform images (Fig. 3A-D). We used the original object and texform images from Long et al. (2018).

We first created a set of original object images (30 each for animate and inanimate images) to confirm that the monkeys could successfully classify these original images. We then created a texform set (30 each for animate and inanimate; Fig. 3A). The texform images were chosen such that they were not made from the original images shown to the monkeys because the monkeys might have remembered the original images. After testing the generalization performance of this texform set, we trained the monkeys on this texform classification for 7-8 sessions (Fig. 3C). Finally, we created another texform set (30 each for animate and inanimate; Fig. 3D) and tested whether the monkeys could generalize the learned texform classification rule to new texform images. The set contained both big and small objects for animate and inanimate images, and we evenly distributed them for each set.

### Monkey experimental procedures

#### Object drag tasks using grayscale image sets

We performed six tasks using grayscale image sets (Fig. 1B, G). The procedure for each task consisted of training and generalization test sessions. During the training sessions, an image from the training set was pseudo-randomly presented to the monkey in each trial. Training lasted for 4-6 sessions (days) depending on the monkey’s learning speed. In the early training phase of each task, we had a portion of trials in which we showed only the correct target box until the monkey performance reached above *∼* 70%. These trials were used to maintain the monkey’s engagement and were excluded from the data analysis. Immediately after the training sessions, we had a generalization session, in which we started to show generalization images in 50% of the trials. We analyzed only the trials in which these generalization images were presented for the first time.

#### Object drag tasks using large-scale THINGS image sets

We performed three tasks using large-scale THINGS image sets. For each task, we first confirmed that the monkeys performed well on the same task with the grayscale image set. We then started to introduce 100 training images from the THINGS set (Fig. 2A). After we confirmed that the monkey performance was stable for the 100 training images, we gradually introduced generalization images. Each generalization image was shown only once. We controlled the proportion of trials in which generalization images were presented, gradually increasing it up to 50%. In the other 50% of the trials, the training images were repeatedly shown to ensure that the monkeys continued to follow the same rule.

#### More abstract classification tasks

Experiments on more classification tasks (Fig. 4) also largely followed the procedure for the other tasks described above (Fig. 1B). Training for each task continued until the monkey performance plateaued but was terminated after six days regardless of their final performance to fairly compare their performance across tasks with similar training durations. The average performances for the outdoor vs. indoor, fire- vs. water-related, and Western vs. Eastern-culture tasks on the last day of training were 91%, 77%, and 88% correct, respectively. The big vs. small task had two versions: one using the THINGS database and one using the images from Konkle and Oliva (2012). We started with the THINGS set (final performance during training: 94%) and, after its generalization test, switched to training on the Konkle set (final performance during training: 93%).

#### Random stimulus-response association tasks

We conducted two versions of control experiments in which monkeys were trained to associate random object images with targets to check how the absence of a common conceptual rule impaired their performance. In one version (Fig. 3E), the associations of images and targets were randomized such that even images of the same object category could be associated with different targets. In the second version (Fig. 3F), two targets were associated with random concrete object concepts (e.g., “deer”, “banana”; we refer to them as “concrete categories” here to avoid confusion), but images in each concrete category were associated with the same target. Thus, if monkeys remembered the association of concrete categories and targets, they could solve the task and generalize the rule to novel stimuli selected from the same concrete categories.

Both versions of the task used the same number of grayscale object images as the animate vs. inanimate task (96 images in the training set) and followed the same training procedure. The training image set for the second version (Fig. 3F) contained 16 concrete categories for each target (2-6 images per concrete category, 48 images per target). As in the other grayscale object tasks (Fig. 1), these concrete categories were sampled from humans, monkeys, non-primate mammals, reptiles/amphibians, food, plants, tools/everyday objects, and electronics. We also constructed a generalization image set with the same number of images and object categories. All the images used in these control experiments differed from those used in the main tasks.

### Human experimental procedures

#### Object drag tasks using grayscale image sets

We performed the six grayscale image tasks (Fig. 1B, G) with human participants (Fig. 6). Seven participants were assigned to each task. In each trial, an image from the training image set was randomly presented, and the participants had to choose a target as in the monkey experiments. The participants were not informed of the categorization rule and had to infer it from the feedback, but they quickly understood the rule in all tasks (Fig. 6A). Then, each participant continued the task for *∼*1,500 trials, resulting in 15-25 repetitions per image. We also conducted the animate vs. inanimate task using three control image sets (cartoons, silhouettes, outlines; Fig. 6G, H). Four participants were recruited for each set, and each participant performed *∼*1,500 trials. In total, 83,416 trials were collected across all tasks and participants.

#### Object drag tasks with generalization images

We also measured the generalization performance of the human participants to novel stimuli in the various tasks we tested with the monkeys (15 tasks in total; Fig. 4, 7). Nine participants were assigned to each task. For each task, we first asked the participants to categorize the training image set. We continued training until the participants reached 90% accuracy (20-trial sliding window) and viewed all the training images (65-183 trials). We then showed the generalization images only once. The same training and generalization images were used as those in the monkey experiments. In total, 13,754 trials for the generalization images were collected across all tasks and participants.

### Data analysis

#### Choice accuracy

To plot accuracy as a function of the number of repeated stimulus presentations in Fig. 1D inset, we computed the correct rate using all stimuli in the two-target trials at a given number of repetitions across training sessions. The correct rate for each image (Fig. 1E) was calculated using the sessions after the monkey performance reached around 75% correct (the last 3-5 sessions). The generalization performances for novel images and manipulated images were all calculated using only the first trial of each stimulus.

### Model comparison

#### Category classification and behavioral fitting procedures

We evaluated the performance of various vision models in classifying object concepts and fitting monkey behavior by constructing linear classifiers based on these model outputs (Fig. 5, Supplementary Figs. 4, 5). The outputs of many models are highly overparameterized (10^2^ *−* 10^5^ parameters), whereas the number of images we used in each task was limited (10^2^ *−* 10^4^). Therefore, we first performed a dimension reduction on the model outputs and then employed a cross-validation approach to evaluate their performance. The cross validation was performed by constructing a classifier using the training set of each task and calculating the model’s correct rate using the generalization set, mirroring the monkey experimental procedure. For dimension reduction, we performed principal component analysis (PCA) on the model outputs of the training set and selected the first 20 PCs. The number of dimensions did not qualitatively affect our conclusions. We then normalized each dimension and constructed a linear classifier of object concepts or monkey choices using a logistic regression. Finally, we projected the model outputs for the generalization set on the same PC space and calculated the proportion of stimuli correctly classified by the trained linear classifier.

A linear classifier was constructed by fitting the training data to the following linear logistic function:

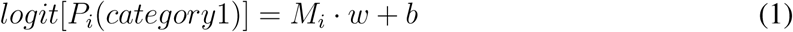

where *P_i_*(*category*1) is the probability of classifying image *i* into category 1, *M_i_* is the dimension-reduced model output for image *i*, *w* is the linear weight, and *b* is a bias parameter. We then computed *P_i_*(*category*1) for each of the generalization images and calculated the correct rate assuming that the model chooses category 1 if *P_i_*(*category*1) is greater than 0.5. For the fitting to monkey behavior, we again performed the linear logistic regression but estimated *w* and *b* to maximize the fit performance to monkey choices in the training trials of the sessions after reaching a plateau performance. The fit performance was quantified as the proportion of generalization images for which the monkey and model choices matched. In the figures, we showed the fit results aggregated across the three monkeys.

We confirmed that changing the details of this model evaluation procedure did not qualitatively affect our results. First, we tested different numbers of dimensions being reduced by PCA (Supplementary Fig. 4A). All the models quickly improved their cross-validated performance as the number of dimensions increased, and their performance reached a plateau before or near 20 dimensions. Second, we tested another dimension reduction technique. We constructed a representational dissimilarity matrix of the images by computing 1 *−* Pearson’s correlation of the model outputs for each image pair. We then performed a classical multi-dimensional scaling (MDS) to obtain a 20-dimensional space capturing the similarity pattern. The classification based on these 20 dimensions was similar to the main results (Supplementary Fig. 4B). Third, we replaced logistic regression with a linear support vector machine (SVM) and found that the classification accuracy did not substantially change (Supplementary Fig. 4C).

#### Model details

##### V1-like model

We followed Pinto et al. (2008) to create a population of V1-like responses, which were generated from a set of Gabor filters of different orientations and spatial frequencies with normalization and non-linear thresholding. In brief, we used Gabor filters (43 *×* 43 pixels) spanning 16 orientations and six spatial frequencies (1/2, 1/3, 1/4, 1/6, 1/11, 1/18 cycles/pixel) with a fixed Gaussian envelope (standard deviation of 9 pixels in both directions) and fixed phase (0) (96 filters in total).

##### Luminance+color model

The model computes marginal statistics of color and luminance information at both the global and local levels. Each pixel intensity of an input image was converted into luminance (*cd/m*^2^) and CIE-u’v’ coordinates via the calibration data of the touchscreen monitors obtained using a spectrometer (X-Rite i1 Pro, X-Rite, Grand Rapids, MI, USA). At the global level, the mean, standard deviation, skewness, and kurtosis of the distributions of luminance, u’, and v’ were calculated for the entire image. At the local level, we segmented the image into 4 *×* 4 blocks and calculated these statistics for each segment. All of these values were normalized and then concatenated as the output of the model.

##### Gist+SF model

The Gist model, adopted from Oliva and Torralba (2006), captures the spatial structure of an image by computing the orientation and spatial-frequency powers of each local image region. Although the extracted features are similar to those of the V1-like model, this model captures larger-scale image structures with more compact representations. In brief, each image was divided into 4 *×* 4 segments and oriented Gabor filters (8 orientations) were applied over different scales (4 scales) in each segment. The average filter energy was subsequently calculated for each filter and segment, and all outputs were concatenated. To capture global image patterns, we also calculated the spatial frequency and orientation power (4 frequencies and 8 orientations) of the whole image. The images were converted to grayscale before the calculation and all the statistics were normalized.

##### Texture model

The texture model consists of statistics that capture the correlation structures of V1-level filter outputs across different scales, orientations, and positions (Portilla and Simoncelli 2000). These statistics explain part of the neural responses in macaque V2 and V4 (Freeman et al. 2013; Kim et al. 2019; Okazawa et al. 2015). We adopted the code of Portilla and Simoncelli (2000), which used steerable pyramids to compute the outputs of oriented bandpass filters. We chose 4 orientations, 4 scales, and 7 spatial neighborhoods to compute spatial correlations. The texture model also included marginal statistics of the luminance distribution and the powers of the V1-level filter outputs.

##### Contour model

For the background-cropped image sets, we computed statistics derived from the outline contour of an object image (Supplementary Fig. 5). The statistics included the aspect ratio (horizontal vs. vertical extent of the contour), the angle of the longitudinal axis (the most elongated axis), the extent of elongation (the ratio between the most elongated axis and its orthogonal axis), the distance of the contour from the image origin at each 5-degree angle, the standard deviation of these distances, and the spatial frequency powers of the contour shape at 4 scales. Thus, these parameters quantified information such as how much and in which direction the object contour is elongated and how smooth the contour is, as well as the coarsely sampled location of the contour itself.

##### Deep neural network (DNN) models

We used seven DNN models commonly used in vision science research —AlexNet (Krizhevsky et al. 2012), VGG16 (Simonyan and Zisserman 2014), ResNet-50 (He et al. 2015), Vision Transformer (ViT-B/32; Dosovitskiy et al. 2020), DINO (Caron et al. 2021), CLIP (Radford et al. 2021) and SigLIP2 (Tschannen et al. 2025). AlexNet, VGG16, and ResNet-50 are convolutional neural networks (CNNs) with different numbers of convolutional layers and network architectures. We used these networks pre-trained on 1,000 object categories in the ImageNet1k dataset. Using the hook mechanism in PyTorch (version 2.0.1), we extracted the activity of the penultimate layer for each image as model outputs. Vision Transformer was used to demonstrate how a DNN with a very different architecture from CNNs performs in our tasks. We extracted the activity of the “encoder” block of ViT-B/32 for each image. DINO is a self-supervised vision transformer that learns visual features from unlabeled images. We chose DINO with ViT-B/16 as the architecture. CLIP and SigLIP2 are recent DNNs that learn visual representations by aligning feature spaces between image and text data (Radford et al. 2021; Tschannen et al. 2025). We chose CLIP and SigLIP2 with ViT-B/32 as the architecture of visual processing and extracted the activity of their visual encoders for each image.

#### Classification based on ventral-pathway neural responses

We also tested whether neural responses recorded from ventral-pathway visual areas could classify the categories in our tasks by using the THINGS ventral stream spiking dataset (TVSD; Papale et al. 2025). These neural data were obtained from two macaque monkeys viewing images from the same THINGS database we used, with a large number of microelectrodes implanted in V1, V4, and inferotemporal (IT) cortex. The dataset includes normalized multi-unit activity (MUA) averaged in a time window centered on the peak response of each region (25-125 ms for V1, 50-150 ms for V4, and 75-175 ms for IT). We used this normalized MUA data to assess how well neural populations (V1: 1024 units, V4: 448 units, IT: 576 units) could classify object categories, following the same procedure used for the model outputs: training a linear classifier on the training image set and testing its performance on the generalization image set (Fig. 5D). To further probe the full decoding capacity of these population responses, we also estimated classification accuracy using 5-fold cross-validation within the generalization sets, which provided more training samples for the classifier (Fig. 5E). Accuracy was evaluated as a function of the number of units by randomly subsampling units from the dataset across 10 permutations (Rajalingham et al. 2020). For the number of units exceeding 20, dimensionality was reduced to 20 using PCA, following the same procedure used for the model classification.

#### Typicality analysis

To test whether monkeys made more errors for less typical images in the dataset, we defined the typicality of each image for each task using background-cropped image sets (Fig. 6I-K) and THINGS image sets (Supplementary Fig. 8), following Kramer et al. (2023). In brief, we calculated Pearson correlations between DNN outputs for each pair of images and averaged the correlations of image pairs within the correct and incorrect category in each task (e.g., average correlations between a dog image and all other animate and inanimate stimuli in the animate vs. inanimate task). Image typicality within each task was defined as the difference in the averaged correlations between the correct and incorrect categories. We used AlexNet outputs in Fig. 6I-J as an example, but other visual DNNs yielded similar results. If an image was atypical for its category, this image typicality should be low. In Fig. 6I-K (background-cropped image set), the typicality of each image was correlated directly with category sensitivity averaged across the three monkeys. In Supplementary Fig. 8 (THINGS image set), the typicality of individual images was first averaged within each concrete object category (e.g., dog, cat), and the resulting category-level typicality was correlated with the monkeys’ category-level performance, averaged across the three monkeys.

### The drift-diffusion model (DDM) fit

To quantify the difficulty of categorizing each stimulus, we used DDM, which can jointly fit choices and RTs (Fig. 6C). The model assumes that the drift rate (category sensitivity) varies across images and reflects stimulus difficulty, whereas the decision bound and non-decision time are stable across images for each participant and task. For each image, the drift-diffusion process forms the decision variable at time *t* (*v*(*t*)) as

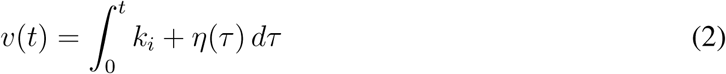

where *k_i_* is the drift rate (category sensitivity) for image *i*. *η*(*τ* ) represents a noise term following a Gaussian distribution with a mean of 0 and an SD of 1. When *v*(*t*) reaches a positive (+*B*) or negative (*−B*) bound, the model commits to the decision associated with the bound (category 1 or 2). The reaction times are modeled as the time required to reach a bound plus an additional delay (non-decision time) common to all images. We modeled the non-decision time as a Gaussian distribution with a mean of *T*_0_ and an SD of *T*_0_*/*3.

The model had two free parameters (*B*, *T* 0) common to all images and the drift rate (category sensitivity; *k_i_*in Eq. 2) as a free parameter for each image (total *n* + 2 free parameters; *n* is the number of images). Higher drift rates indicate that images were easier to classify. We fitted the model to each participant’s choices and RT distributions in each task using maximum likelihood estimation (Luo et al. 2025; Zheng et al. 2025). Given a set of parameters, the RT distributions for the two choices were calculated using the aforementioned model formulation by numerically solving the Fokker-Planck equation. From these distributions, we derived the likelihood of observing the participants’ choices and RTs. Summing the log-likelihoods over all trials yielded the total log-likelihood of the parameter set. We used a simplex search method (*fminsearch* in MATLAB) to determine the parameter set that maximized the summed log-likelihood. As shown in Supplementary Fig. 7, our model accurately fit both monkey and human RTs and correct rates, thereby justifying the use of the drift rate as a metric of stimulus difficulty. Figure 6E-H show the drift rate averaged across participants.

### Sensitivity correlations across individuals

To quantify the similarity of performances within and across species, we calculated Pearson correlations of category sensitivity (k) in Fig. 6L between all pairs of subjects (both monkey and human) for each task. Prior to correlating, k was sign-adjusted per image so that a positive k indicated a bias toward the image’s correct category regardless of category label. Pairwise correlations were averaged within each group (monkey-monkey, human-human, and monkey-human) to yield within- and across-species correlations, which were also averaged across tasks.

### Dissimilarity matrix

We used the generalization performance of images across all tasks to construct the dissimilarity matrix among humans, monkeys, and models (Fig. 7A). As in Fig. 5C, we focused on the first trial of each generalization image and calculated the average correct rate across subjects for humans and monkeys. For the models, we calculated the probability of correct classification for the same image based on the likelihood derived from a logistic classifier trained on the training images. We then concatenated all image performances across tasks. The dissimilarity was defined as Euclidean distance; thus, both the overall correct rates and the patterns of performance across images influenced the values. Using 1*−* Pearson’s correlation yielded similar results, but low- and mid-level models became more distant from each other because they exhibited distinct error patterns. t-SNE (Fig. 7C) also used Euclidean distance, and its perplexity parameter was fixed to 4.

## Supplementary figures

**Figure S1:**
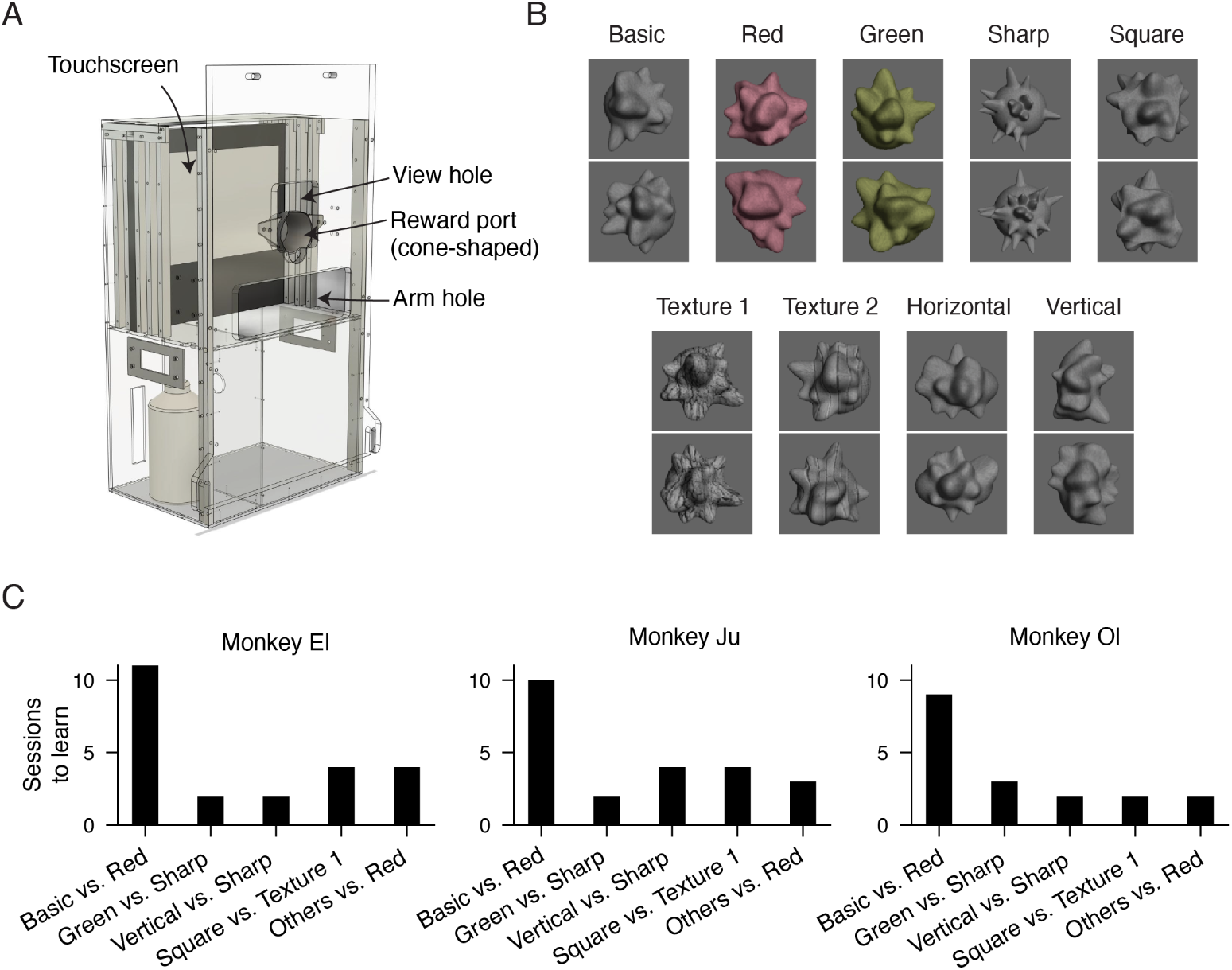
Pretraining of monkeys on the task structure using shape stimuli. (**A**) Behavioral testing system attached to the monkeys’ home cage. Monkeys reached for a touchscreen through an arm hole in the front panel and received a juice reward from a cone-shaped reward socket (Kawaguchi et al. 2019). While performing tasks, they placed their mouth on the reward socket and looked at the screen through a viewing hole. (**B**) Prior to the main concept classification tasks, we trained the monkeys to perform the classification of amorphous shape stimuli. The task sequence followed the main design (Fig. 1A), but the monkeys had to categorize shape stimuli according to their shape, color, or texture. We created nine groups of stimuli as shown in this panel and chose two of them to classify for each task. In each group, there were 50 images with random bump positions. (**C**) The number of sessions (days) required for the monkeys to learn each shape classification task plotted in the trained order. Prior to the first task, monkeys were trained to touch a shape stimulus and move it to a target box. Then, in the first “Basic” vs. “Red” task, they were presented with two target boxes and had to learn to select one of them based on a stimulus feature. This initial stimulus-response association training took 9-11 days. The monkeys were subsequently trained to classify different pairs of shapes in 2-5 days each. In the “Others” vs. “Red” task, the monkeys had to choose one target for red stimuli and the other target for stimuli randomly selected from the remaining eight stimulus categories.

**Figure S2:**
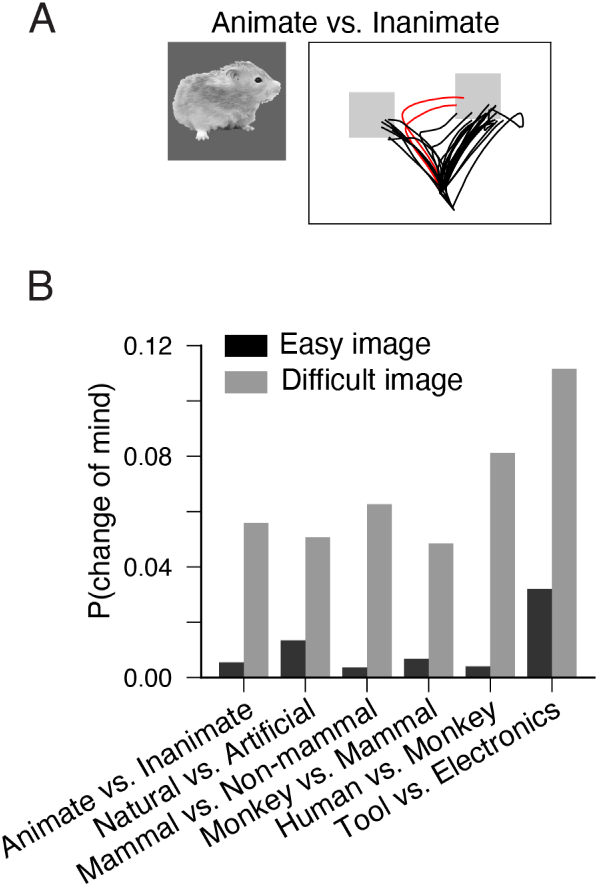
Monkey reach trajectories reflected their decision-making process. (**A**) Monkeys moved a stimulus to a target with a straight path in most trials, but they occasionally made detours as if they changed their mind (red lines; Kaufman et al. 2015). These trials were detected by looking for trajectories whose average position was the opposite to the position of the chosen target with respect to the midline. The right target box corresponded to the animate category in this example. (**B**) Monkeys exhibited these detour trajectories much more often for difficult images. For each task, we selected the 10 easiest and most difficult images based on correct rates and plotted the probability of trials in which the monkeys exhibited detour trajectories. The plots aggregate the data from the three monkeys.

**Figure S3:**
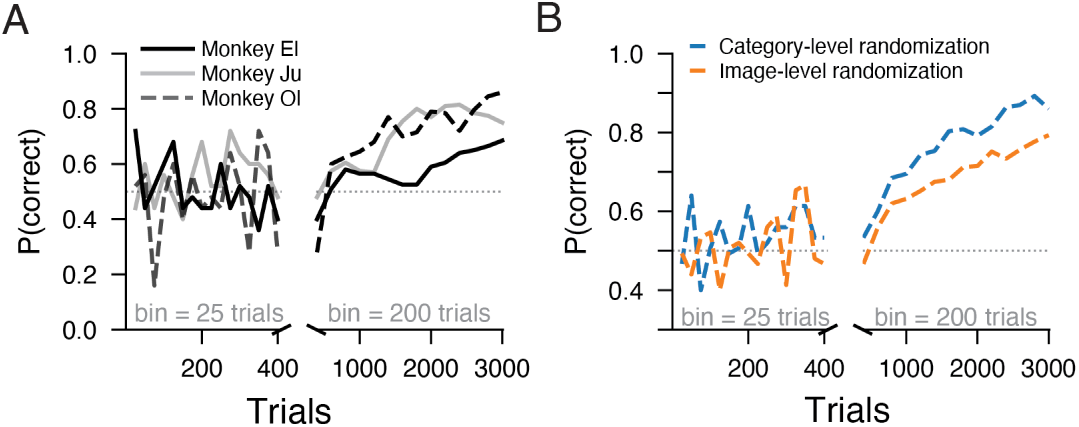
Trial-level learning time courses of control tasks. Learning time courses shown in Fig. 3C and 3G are replotted as a function of trial count. Conventions are the same as in Fig. 1H.

**Figure S4:**
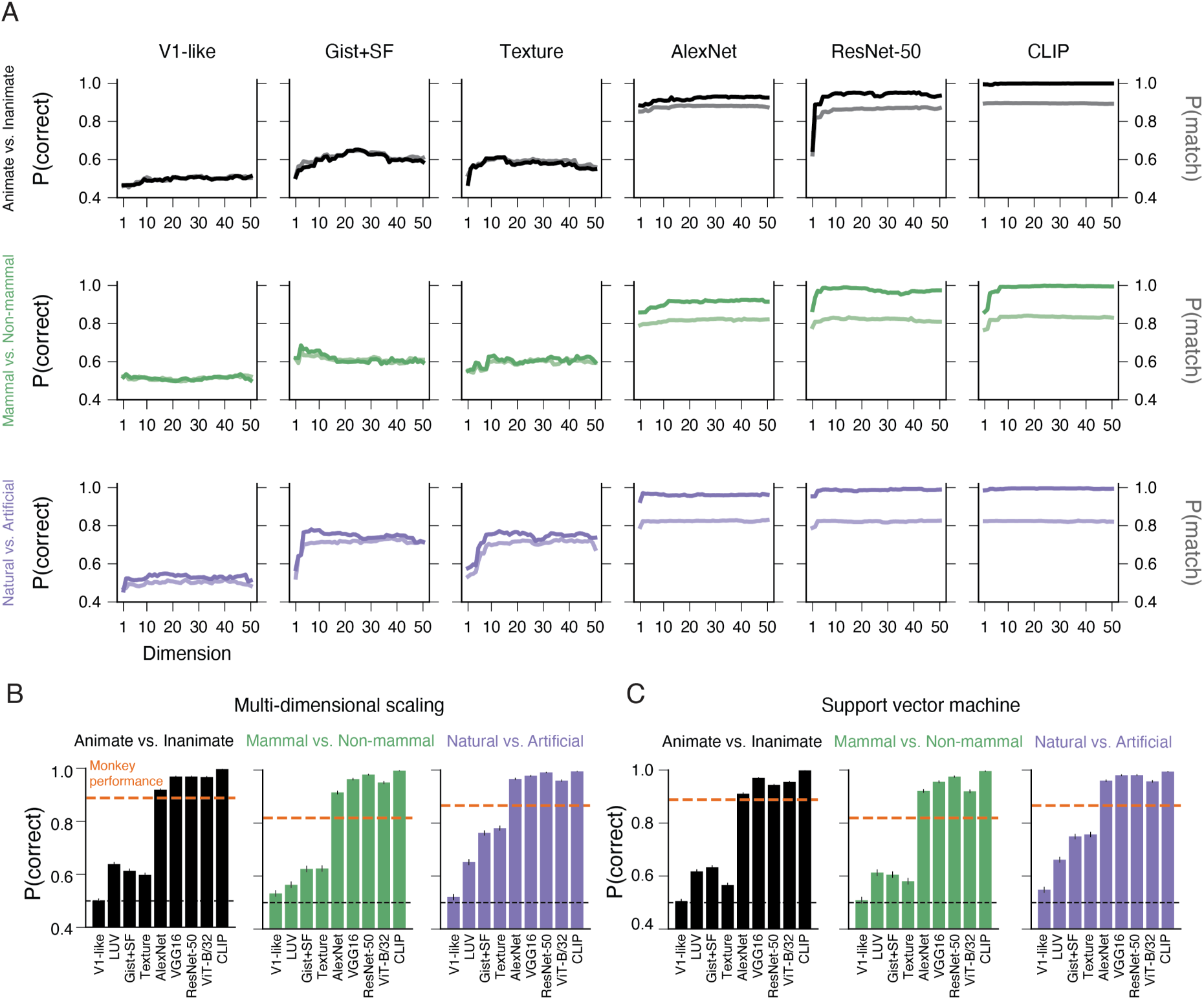
Model performance did not depend on classification or dimension reduction methods. (**A**) We confirmed that the classification and behavioral fitting performances of the models (Fig. 5) did not strongly depend on the number of dimensions of the model parameters reduced by principal component analysis (PCA). In each panel, the darker line represents the model performance in correctly classifying object concepts, and the lighter line represents the model performance in fitting monkey choices. In both cases, the fitting was performed using the training image sets, and the performance was evaluated using the generalization image sets used in the monkey experiments shown in Fig. 2. (**B, C**) Model performance was not strongly dependent on the methods used for dimension reduction and classification. We tested multi-dimensional scaling for dimension reduction (B) and used a support vector machine instead of logistic regression for classification (C) and found that the results did not change qualitatively.

**Figure S5:**
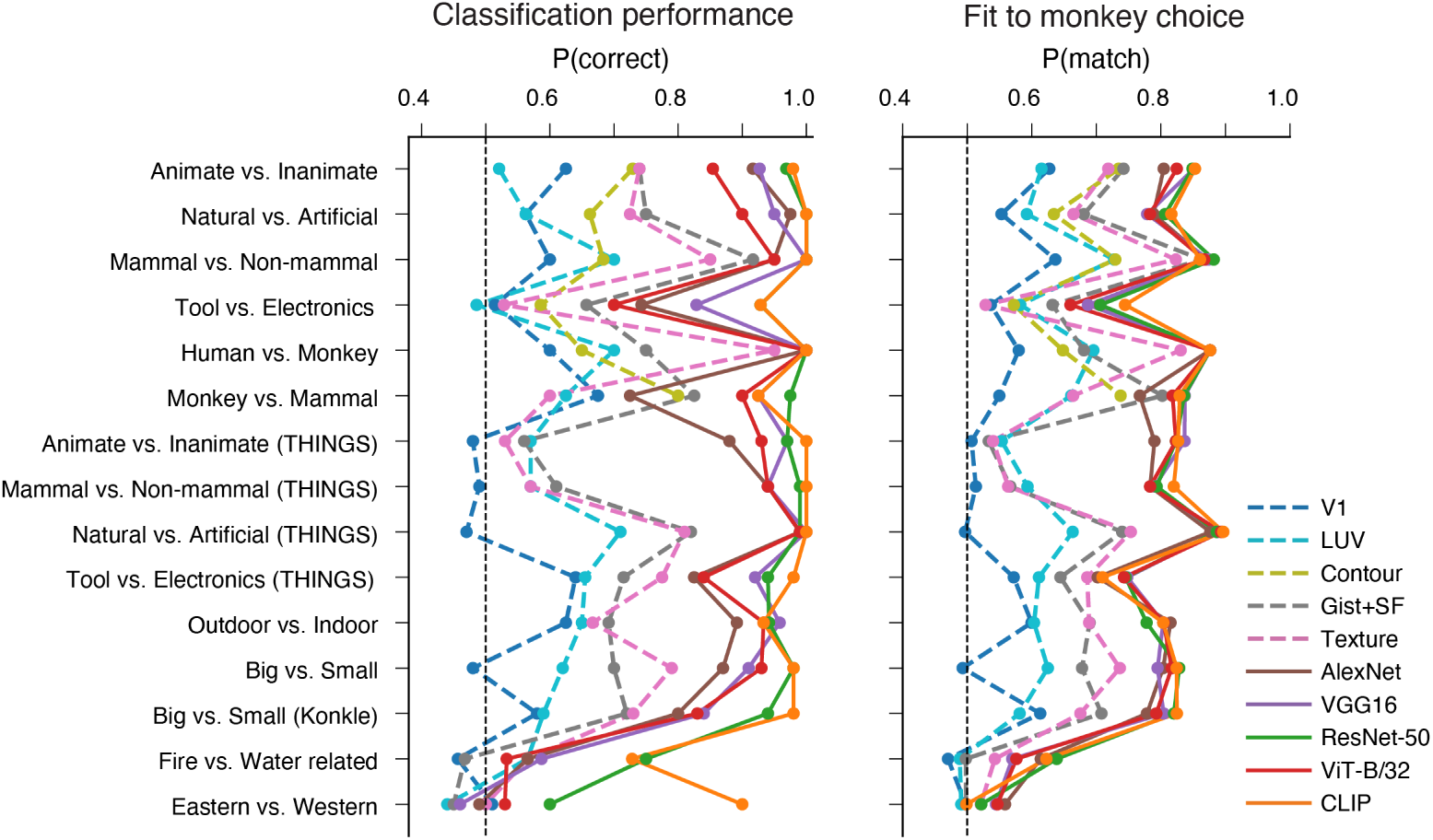
Model classification and behavioral fitting performance for all tasks. Summary of model performance for all tasks shown in Figs. 1, 2, and 4. For the tasks using large-scale THINGS images (Fig. 2), we used only the first 100 generalization stimuli to compute the performance for proper comparison; thus, the performances do not exactly match those shown in Fig. 5A. The classification performances of the models shown in Fig. 5C are the same.

**Figure S6:**
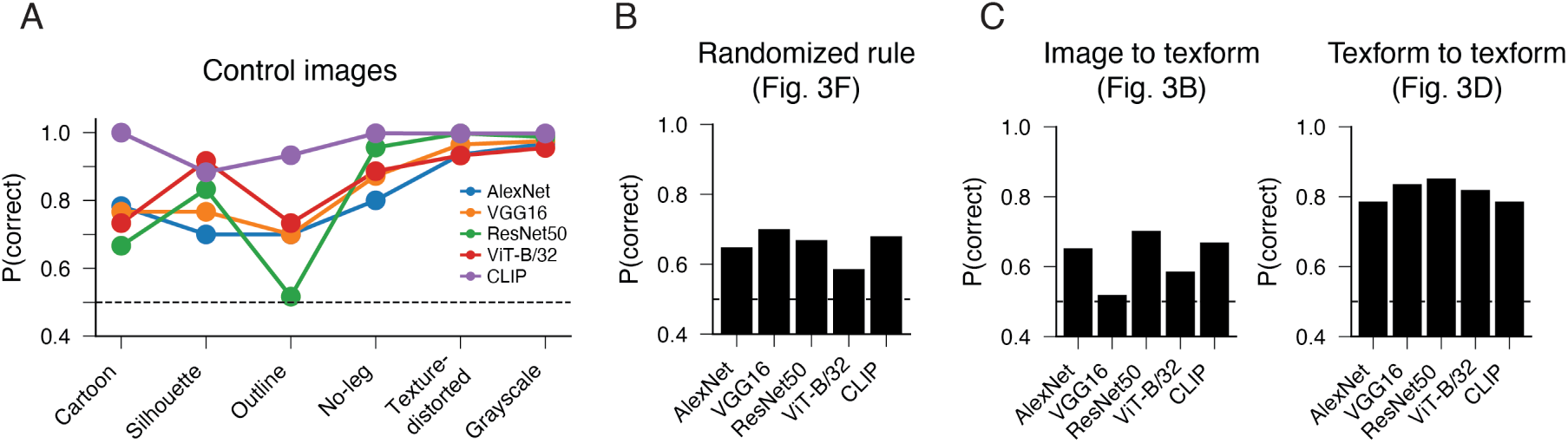
Model classification performance for control tasks. (**A**) DNNs could classify modified stimuli outside the range of natural images (cartoon, silhouette, outline images) and other control images with reasonable accuracy, consistent with monkeys (Fig. 2D, 6G). The performance tended to be lower for the outline images, which is consistent with the monkey and human behavioral results (Fig. 6G). (**B**) DNN performance on the rule randomization task (Fig. 3F), in which the same concrete object categories (e.g., human, flower) were associated with the same targets but there was no common rule for classifying all images. Here, linear classifiers trained on DNN outputs failed to generalize the individual association rules to novel images, similar to monkey performance (Fig. 3H). (**C**) DNN performance on the texform classification task (Fig. 3A-D). As in the monkey experiments, DNN classifiers were first trained on the animate vs. inanimate task and then tested on the texform version of the animate vs. inanimate images, on which they performed poorly (left). The classifiers were then trained on the texform set itself and tested on a separate texform set, on which they performed better (right). These patterns were qualitatively similar to the monkeys’ behavior.

**Figure S7:**
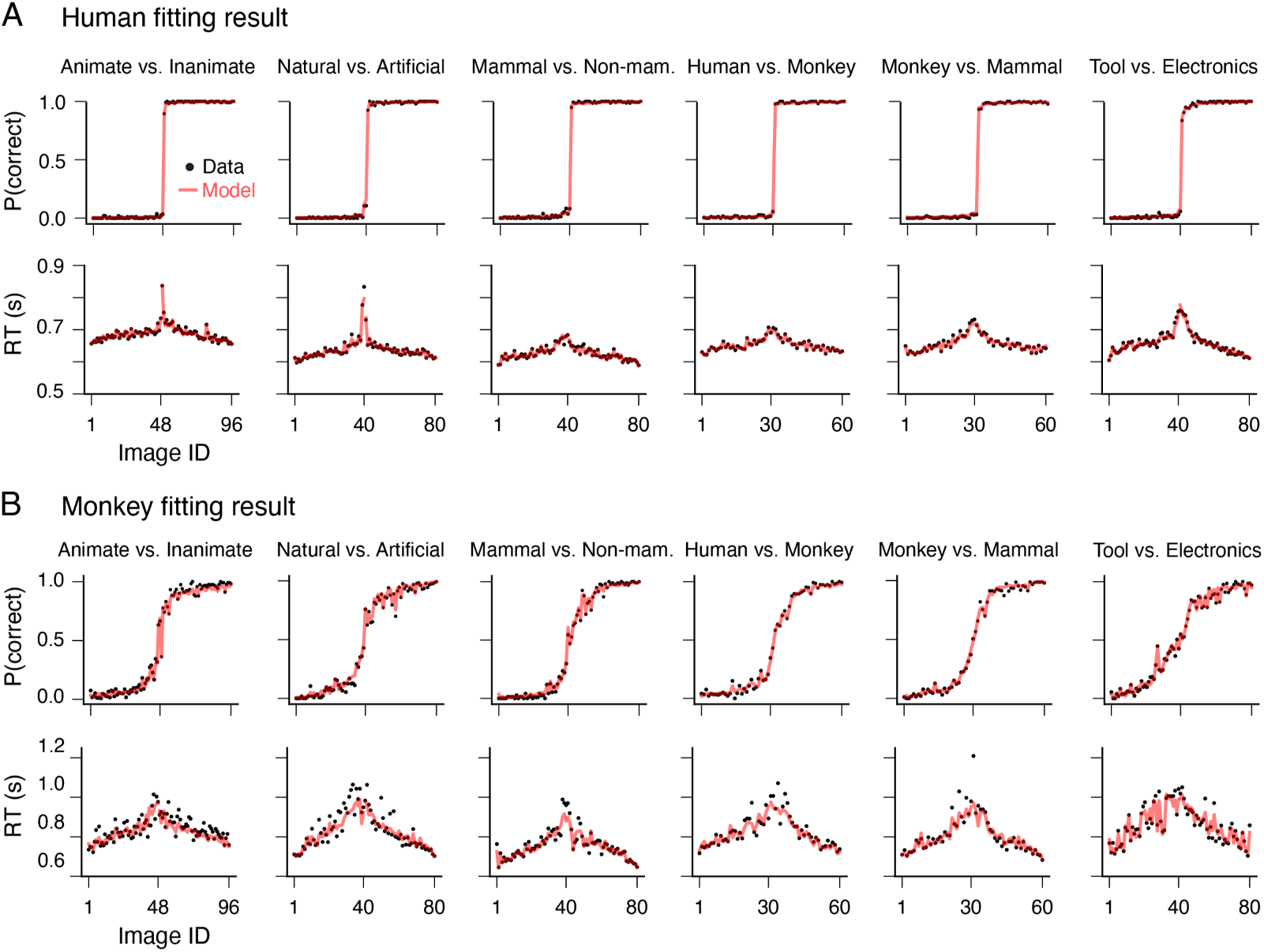
Drift diffusion model (DDM) could accurately fit choices and reaction times. (**A, B**) Fitting of the DDM (Fig. 6C) to the behavioral data of individual tasks for humans (A) and monkeys (B). The conventions are the same as those in Fig. 6D. Model fit was performed for the tasks using grayscale image sets (Fig. 1), in which the monkeys repeated many trials for the same stimuli. See Methods for the details of the model fit. The image IDs were sorted by the sensitivity parameter (k_i_ in Eq. 2) of the model fit averaged across participants for each task. The fit lines are not smooth because the plots are the average across participants who had different rank orders of sensitivity across image IDs.

**Figure S8:**
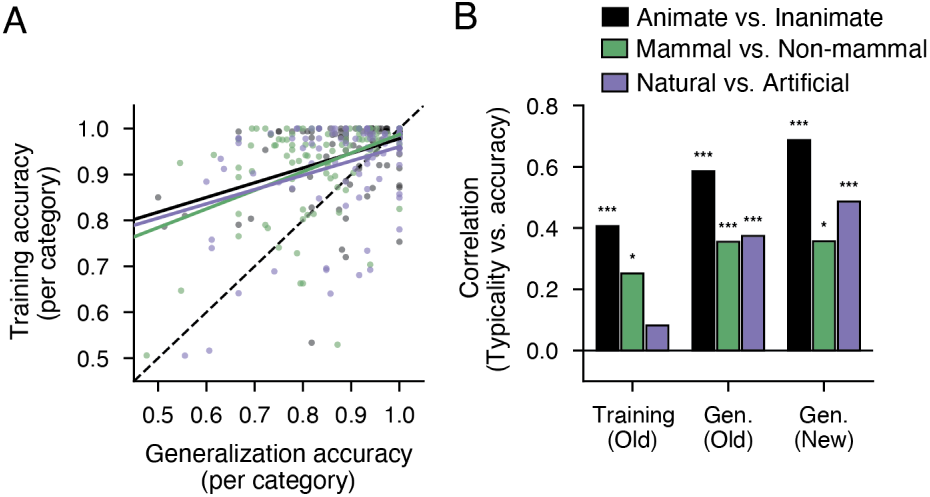
The effect of repeated training and image typicality on monkeys’ generalization performance. (**A**) In the THINGS tasks (Fig. 2), we contrasted monkeys’ performance for stimuli repeatedly presented (training set) with performance for new, generalization stimuli drawn from the same categories (“old” in Fig. 2A). To assess how repeated reinforcement of the training set shaped behavior, we plotted the mean performance for each object category (one dot per category) between these two conditions. We still found moderate positive correlations between the conditions (R > 0.38, p < 0.001 across all three tasks), suggesting that the strategies monkeys applied when categorizing generalization stimuli kept influencing the categorization of repeatedly presented stimuli. (**B**) We then tested if the image typicality (Fig. 6I) accounted for monkeys’ correct rates across object categories in the THINGS tasks. We calculated typicality for each image (see Methods), averaged it across the images for each object category, and compared it with the monkeys’ average correct rate for that category. For all cases except for the training set in the natural vs. artificial task, we found significant correlations, indicating that typicality influenced the monkeys’ performance. ^∗^p < 0.05, ^∗∗∗^p < 0.001.

Movies 1-3 Example videos showing the monkeys’ performance on the tasks using the large-scale THINGS database (Fig. 2; Movie 1: monkey Ju performing the animate vs. inanimate task, Movie 2: monkey El performing the natural vs. artificial task, Movie 3: monkey Ol performing the mammal vs. non-mammal task). The videos were taken during the generalization session, where half of the images were from the generalization set, which was shown only once in each task. The other half were from the training set. The upper parts of the videos are screenshots of the monitor the monkey was looking at.

